# Pseudomonas aeruginosa secreted protein PA3611 promotes bronchial epithelial cells epithelial-mesenchymal transition through TGF-β1 inducing p38/miRNA/NF-κB pathway

**DOI:** 10.1101/2020.10.14.339044

**Authors:** Lei Shu, Sixia Chen, Xiaolin Chen, Shaoqing Lin, Xingran Du, Kaili Deng, Jing Wei, Yang Cao, Jiaxin Yan, Ziyan Shen, Ganzhu Feng

## Abstract

Pseudomonas aeruginosa (PA) is one of the important pathogens, which has been proven to colonize and cause infection in the respiratory tract of patients with structural lung diseases, and further lead to bronchial fibrosis. Epithelial-Mesenchymal Transition (EMT) of bronchial epithelial cells plays a vital role in the process of bronchial fibrosis. Up to the present, the research on bronchial epithelial cells EMT caused by secreted virulence factors of PA has not been reported. In our present study, we found that PA3611 protein stimulation induced the bronchial epithelial cells EMT with up-regulation of mesenchymal cell markers and down-regulation of epithelial cell markers. Meantime, TGF-β1 secretion was markedly increased, IκBα expression was significantly decreased, and NF-κB p65 subunit phosphorylation was markedly enhanced, in addition, the levels of miR-3065-3p and miR-6802-3p expression and p38 MAPK phosphorylation were obviously increased in bronchial epithelial cells after PA3611 stimulation, further research revealed that PA3611 promoted EMT occur through TGF-β1 induced p38/miRNA/NF-κb pathway. The function of PA3611 was also verified in PA-infected rats and results showed that ΔPA3611 could reduce lung inflammation and EMT. Overall, our results revealed that PA3611 promotes EMT via simulating the production of TGF-β1 induced p38/miRNA/NF-κB pathway-dependent manner, suggesting that PA3611 acts as a crucial virulence factor in bronchial epithelial cells EMT process and has potential use as a target for clinical treatment of bronchial EMT and fibrosis caused by chronic PA infection.

**Author summary:** Structural lung disease can increase the chance of chronic infection, including infected by Pseudomonas aeruginosa, which can cause lung structure damages and affect lung functions in further, and forming a vicious circle of intertwining, ultimately, it leads to pulmonary fibrosis. EMT of bronchial epithelial cells plays a vital role in the process of bronchial fibrosis. However, the relationship and mechanism of PA infection leads to the destruction of lung structure and bronchial epithelial cells EMT are still not very clear. We found pseudomonas aeruginosa secreted protein PA3611 can stimulate bronchial epithelial cells EMT through up-regulation of mesenchymal cell markers α-SMA and Vimentin expression and down-regulation of epithelial cell markers E-cadherin and Zonula Occludens-1. Meantime, TGF-β1 secretion was markedly increased, IκBα expression was significantly decreased, and NF-κB p65 subunit phosphorylation was markedly enhanced, in addition, the levels of miR-3065-3p and miR-6802-3p expression and p38 MAPK phosphorylation were obviously increased in bronchial epithelial cells after PA3611 stimulation, further studies suggested that PA3611 was shown to promote EMT occur through TGF-β1 induced p38/miRNA/NF-Kb pathway. Our results revealed that PA3611 promotes EMT via simulating the production of TGF-β1 induced p38/miRNA/NF-κB pathway-dependent manner, suggesting that PA3611 acts as a crucial virulence factor in bronchial epithelial cells EMT process and as a potential target for the treatment of chronic structural lung diseases.

## Introduction

Pseudomonas aeruginosa (PA) is a kind of non-fermented gram-negative bacteria that can cause various acute and chronic infections, including infections of the blood system, urinary system, central nervous system, bone and joints, etc. It is also one of the most common pathogens that cause hospital-acquired infections including pneumonia[1], meanwhile, an opportunistic pathogen and the main cause of morbidity and death in cystic fibrosis patients and immunocompromised individuals[2]. In recent years, with the continuous increasing clinical isolation rates of multi-drug-resistant and pan-drug-resistant PA, the mortality of patients with PA infection has continued to be at a high level[3], which brings great challenges to clinical treatment and poses a serious threat to human health[4]. Therefore, it has been an urgent and important task to actively seek effective countermeasures against PA infection. PA is widespread in nature, and is prone to translocation and infection of the respiratory tract, especially in patients with structural lung diseases such as Chronic Obstructive Pulmonary Disease (COPD), bronchiectasis, and cystic fibrosis[5,6]. PA infection may promote the formation of bronchial fibrosis such as thickening of the airway wall and lumen stenosis in patients with COPD, bronchiectasis and cystic fibrosis, thereby aggravating the irreversible airflow limitation of patients. Studies have shown that the pathological manifestations of bronchial fibrosis are mainly airway smooth muscle thickening, extracellular matrix deposition, myofibroblast proliferation and airway epithelial cell-mesenchymal transition (Epithelial-Mesenchymal Transition, EMT)[7]. Among them, EMT plays a key role in the process of bronchial fibrosis[8]. As is known to all, EMT is cells lose the original polarity, into mesenchymal cells form and characteristics of pathological process, the expression of original epithelial cell markers, such as E-cadherin (E-cad) and tight junction protein 1 (Zonula Occludens-1, ZO-1) reduced, and mesenchymal cells markers, such as α-Smooth Muscle Actin (α-SMA), N-cadherin (N-cad) and Vimentin increased, and some Matrix Metalloproteinases (MMPs) activities were also increased. Among them, α - SMA, Vimentin and MMP-9 high expression can be used as important signs of cell EMT[9].

In recent years, although EMT studies on airway epithelial cells in patients with COPD have been reported, but most of them were cigarette smoke-induced COPD model as the main research object or only take the total lytic substance of PA bacteria as the cell irritant[10,11,12], however, studies on the EMT of bronchial epithelial cells caused by the secretion of virulence factors under PA infection have not been reported. PA3611 is a toxic protein secreted under PA infection (UniProt ID: Q9HY15). Its encoding gene is 411 base pairs in length (including signal peptide) and its molecular weight is 14KDa. Proteomic analysis showed that PA3611 may be a virulence factor regulated by the quorum sensing system[13]. Some preliminarily studied have shown that the spatial structure of PA3611 is composed of five-chains (B1-B5) and five-helix (H1-H5) domains[14], however, its specific biological functions, especially its related effects on infected host cells, have not been reported.

In this study, we used in vivo and in vitro experiments to clarify the PA3611 protein secreted by PA after infection of bronchial epithelial cells to induce EMT and the relative mechanism, in order to provide new clinical treatment for chronic PA infection leading to bronchial EMT and fibrosis.

## Results

### Heterologous expression and purification of recombinant PA3611

The PA3611 protein was synthesized and purified for further investigation. The PA3611-encoding gene was amplified by PCR, and cloned into the plasmid pET-28a(+), then transformed into *Escherichia coli BL21* (DE3). The positive clones were confirmed by sequencing, the results of which were shown in Supplemental fig 1(A, B). The recombinant protein PA3611 was expressed in *E. coli BL21*(DE3) and was subsequently purified with Ni-IDA resin.

### PA3611 inhibits the proliferation of bronchial epithelial cells and promotes the transformation of epithelial cells into mesenchymal cells

To explore the function of PA3611 in bronchial epithelial cells, a cell counting kit-8 (CCK-8) assay was used to estimate its effect on bronchial epithelial cells proliferation. The results demonstrated that PA3611 infection significantly inhibited 16HBE and RTE cells proliferation, and this effect increased with higher concentration of PA3611 and longer incubation times (Fig. 1A, B), which indicated PA3611 inhibited cell proliferation was in a time and dose-dependent manner.

**Figure 1:**
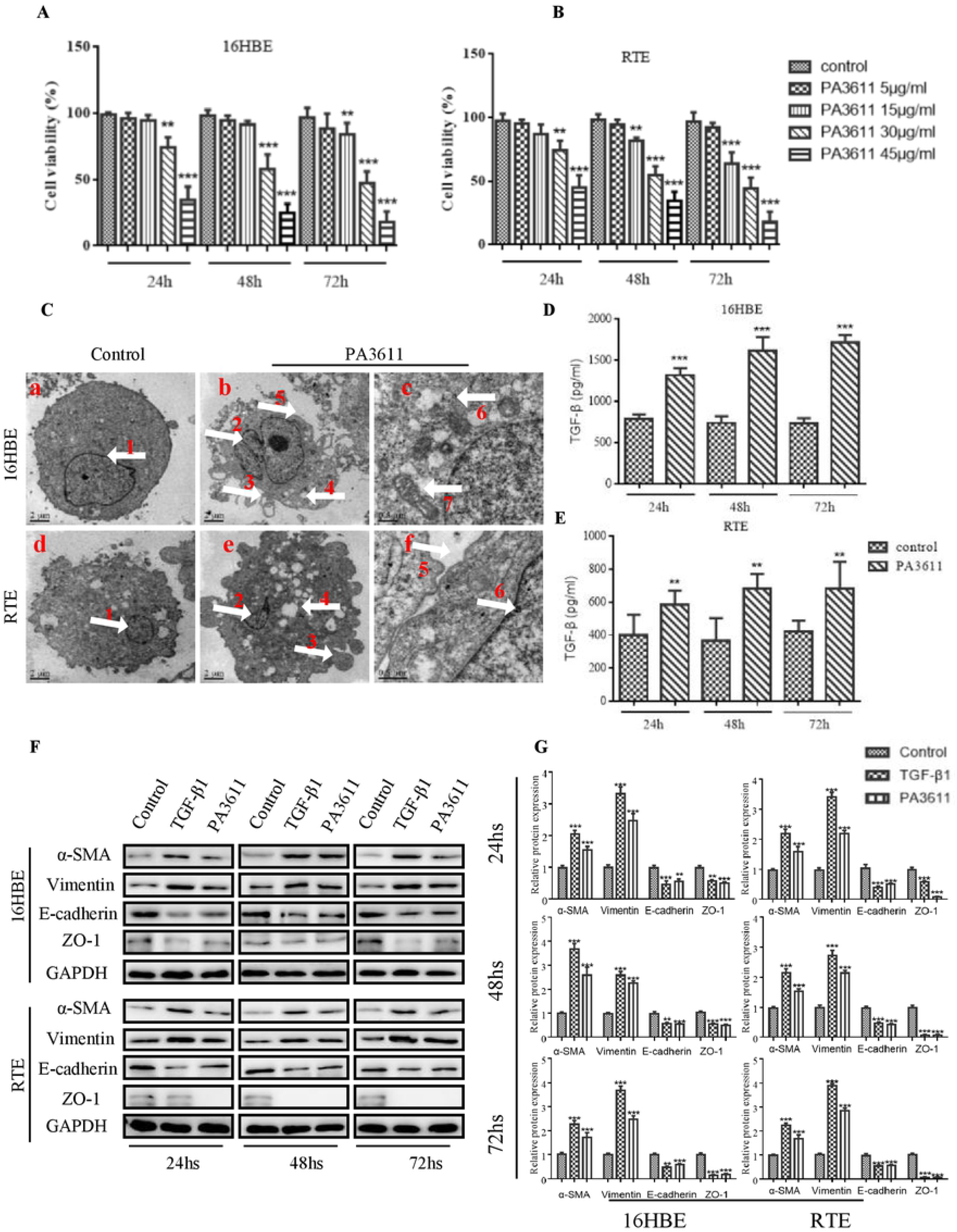
Effects of PA3611 on the proliferation of bronchial epithelial cells and epithelial mesenchymal transition (EMT). 16HBE and RTE cells were treated with PA3611 at the different concentration (5μg, 15μg, 30μg and 45μg) for 24, 48 or 72hs. A CCK-8 assay was performed to estimate cell proliferation (A, B). Transmission electron microscopy was used to observe the ultrastructure of the above cells treated with PA3611 (30μg/ml) for 72 hs (C). TGF-β1 levels in 16HBE and RTE cells culture supernatants were evaluated by enzyme-linked immunosorbent assay treated with PA3611 (30μg/ml) for 24hs, 48hs or 72hs (D, E). EMT-related proteins (α-SMA, Vimentin, E-cadherin and ZO-1) were tested with Western-blot in 16HBE and RTE cells at 24, 48 or 72hs time-points (F and G, PBS as control group, TGF-β1(2 ng/ml), as positive control). All the presented graphs are representative of three independent experiments. Data are presented as the means±SD. *, P <0.05, **, P <0.01, ***, P <0.001, vs. control group.

In order to further understand the in internal ultrastructure changes of the above two bronchial epithelial cells stimulated by PA3611 protein, transmission electron microscopy was used to observe the ultrastructure of the above cells treated with PA3611 (30μg/ml) for 72 hours. The results showed that: the intercellular gaps were widened, intercellular connection disappeared, and filamentous pseudopodia appeared, while the nucleus were heteromorphic and the perinuclear gaps were widened, Mitochondria were swollen, distorted and intracellular glycogen were increased, as shown in Fig. 1C. These results indicated that PA3611 protein could promote the transformation of bronchial epithelial cells into mesenchymal cells under the action of certain concentration and time.

### PA3611 stimulation of TGF-β1 production is accompanied by promotes bronchial epithelial cells epithelial-mesenchymal transition and p65 phosphorylation

TGF-β1 is a key cytokine and participates in the EMT process[15,16], to determine whether PA3611 promoted bronchial epithelial cells EMT by stimulated TGF-β1 production, the TGF-β1 levels in 16HBE and RTE cells culture supernatants were evaluated by enzyme-linked immunosorbent assay at 24hs, 48hs and 72hs times-points (ELISA). The results showed that TGF-β1 levels in supernatants were increased after PA3611 stimulation both in 16HBE and RTE cells at all three time-points (Fig. 1D, 1E). We further examined the effect of PA3611 on the expression of EMT-related marker proteins in above cells, compared with control group (PBS), TGF-β1(2 ng/ml, positive control) and PA3611 (30μg/ml) treated groups both increased α-SMA and Vimentin expression meanwhile decreased E-cadherin ZO-1 proteins expression at 24hs, 48hs and 72hs times-points (Fig. 1F, G), similar results were seen from mRNA levels (Fig. 2A).

**Figure 2:**
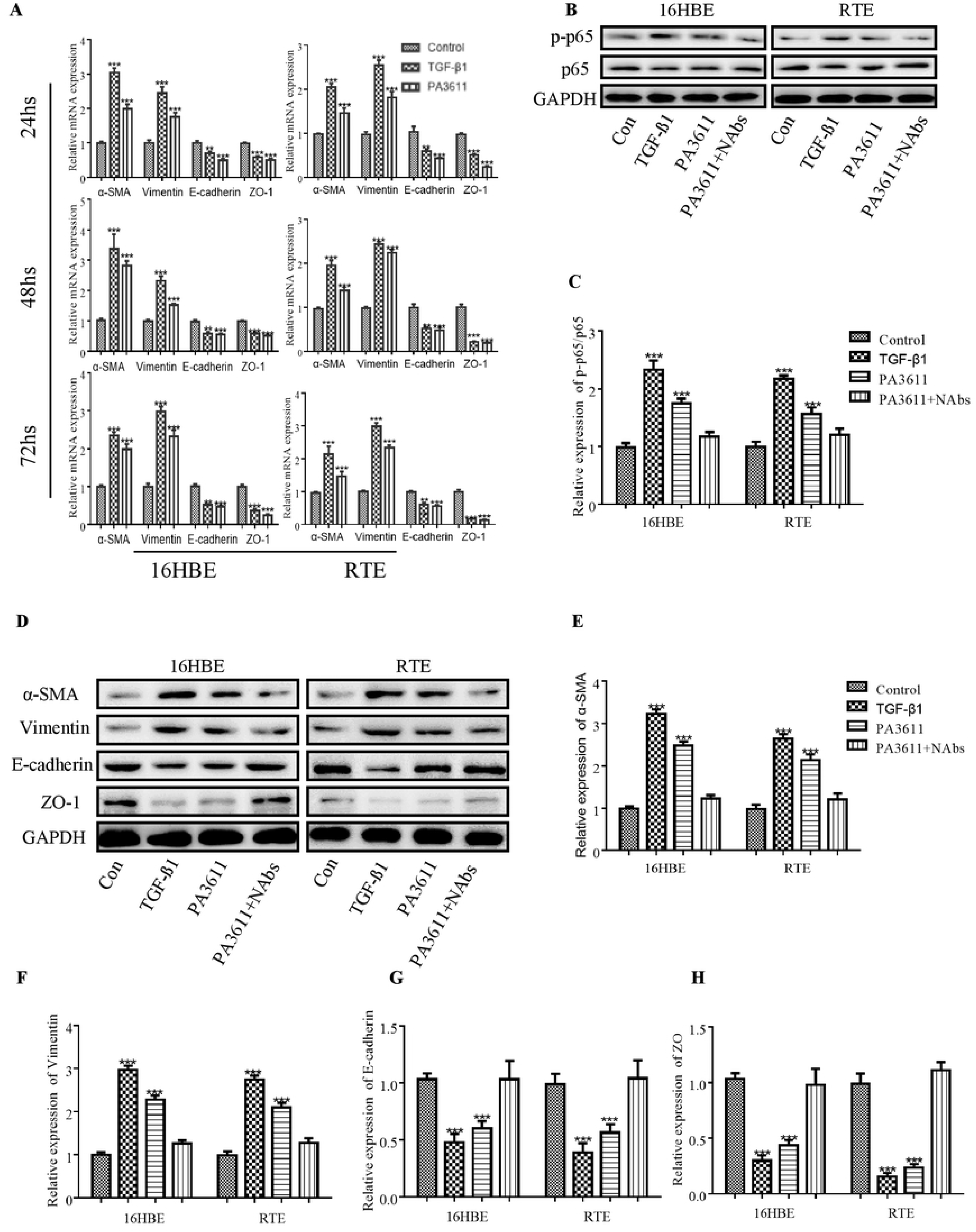
Effects of PA3611 and TGF-β1 on the expression of p65 phosphorylation and EMT relative markers. 16HBE and RTE cells were treated with PA3611(30μg) and TGF-β1(2ng/ml) for 24, 48 or 72hs. EMT-related genes (α-SMA, Vimentin, E-cadherin and ZO-1) were tested with qRT-PCR in 16HBE and RTE cells at 24, 48 or 72hs time-points (A, PBS as control group, TGF-β1(2ng/ml), as positive control). 16HBE and RTE cells were treated with PA3611(30μg), TGF-β1(2ng/ml) or PA3611(30μg) plus TGF-β1 neutralizing antibody(10μg/ml) for 24hs, p65 phosphorylation (B, C) and EMT relative markers (α-SMA, Vimentin, E-cadherin and ZO-1) were tested with Western-blot(D-H). All the presented graphs are representative of three independent experiments. Data are presented as the means±SD. *, P <0.05, **, P <0.01, ***, P <0.001, vs. control group.

A lot of studies have revealed that NF-κB and p65 phosphorylation play a key role in the EMT process[17,18,19,20]. In order to clarify the effect of PA3611 on the expression of p65 p-p65 and EMT process, 16HBE and RTE cells were treated with PBS (negative control), TGF-β1 (2ng/ mL, positive control), PA3611 (30ug/mL), and PA3611 (30ug/mL) + TGF-β1 neutralizing antibody (10μg/mL) for 24 hs, the results indicated that: TGF-β1 and PA3611 increased the expression of p-p65, α-SMA and Vimentin and decreased the expression of E-cadherin and Zo-1 obviously compared with PBS group, while PA3611+ TGF-β1 neutralizing antibody significantly decreased the expression of p-p65, α-SMA and Vimentin, and promoted the expression of E-cadherin and Zo-1, of which the change trend was similar to that in PBS group (Fig 2 B-H). This results indicated that PA3611 promoted EMT via stimulating TGF-β1 production induced phosphorylation of p65.

### PA3611 upregulates miR-3065-5p and miR-6802-3p in PA3611 infected bronchial epithelial cells

Recent studies have found that miRNAs play a specific and important role in regulating related cells EMT[21,22], in order to identify miRNAs associated with the process of PA3611 infected bronchial epithelial cells EMT, a microarray analysis was conducted to miRNAs that were differentially expressed between control and PA3611 infected groups. Differentially expressed miRNAs were identified using a screening criterion of |Fold change|≥2 and identified miRNAs are shown in Supplemental fig 1C. Among these, miR-3065-5p and miR-6802-3p were significantly upregulated in PA3611-treated and evaluated by qRT-PCR. (Suppl Fig. 3A-B).

### p65 upregulates the levels of α-SMA and Vimentin expression without affecting TGF-β1, p38 phosphorylation, IκB αand miRNA expression

To verify the effect of p65 on EMT process and related pathway factors expression, p65 was overexpressed in 16HBE via transfection with pcDNA3.1/p65 cDNA or knocked down through transfection with a specific small interfering RNA (siRNA) directed against p65. The results showed that both p65 overexpression and PA3611 upregulated the expression of α-SMA and Vimentin, and decreased the levels of E-cadherin and Zo-1from gene and protein levels, moreover, the expression levels changed more significantly after treated with PA3611 plus p65 overexpression (Fig 3 E-I; Suppl Fig. 2D-G). However, the TGF-β1, p38, IκBα and miRNAs expression were not affected by either overexpression or knocked down of p65 (Fig 3 A-D, J; Suppl Fig. 2A-C, H-I), after plus PA3611, the expression of TGF-β1, p38 and miRNAs were significantly up-regulated and IκBα down-regulated regardless of p65 overexpression or knocked down (Fig 3 A-D, J; Suppl Fig. 2A-C, H-I). These results indicated that PA3611, not p65 has a regulatory effect on secretion of TGF-β1, phosphorylation of p38 and expression of IκBα, miR-3065-5p and miR-6802-3p.

**Figure 3:**
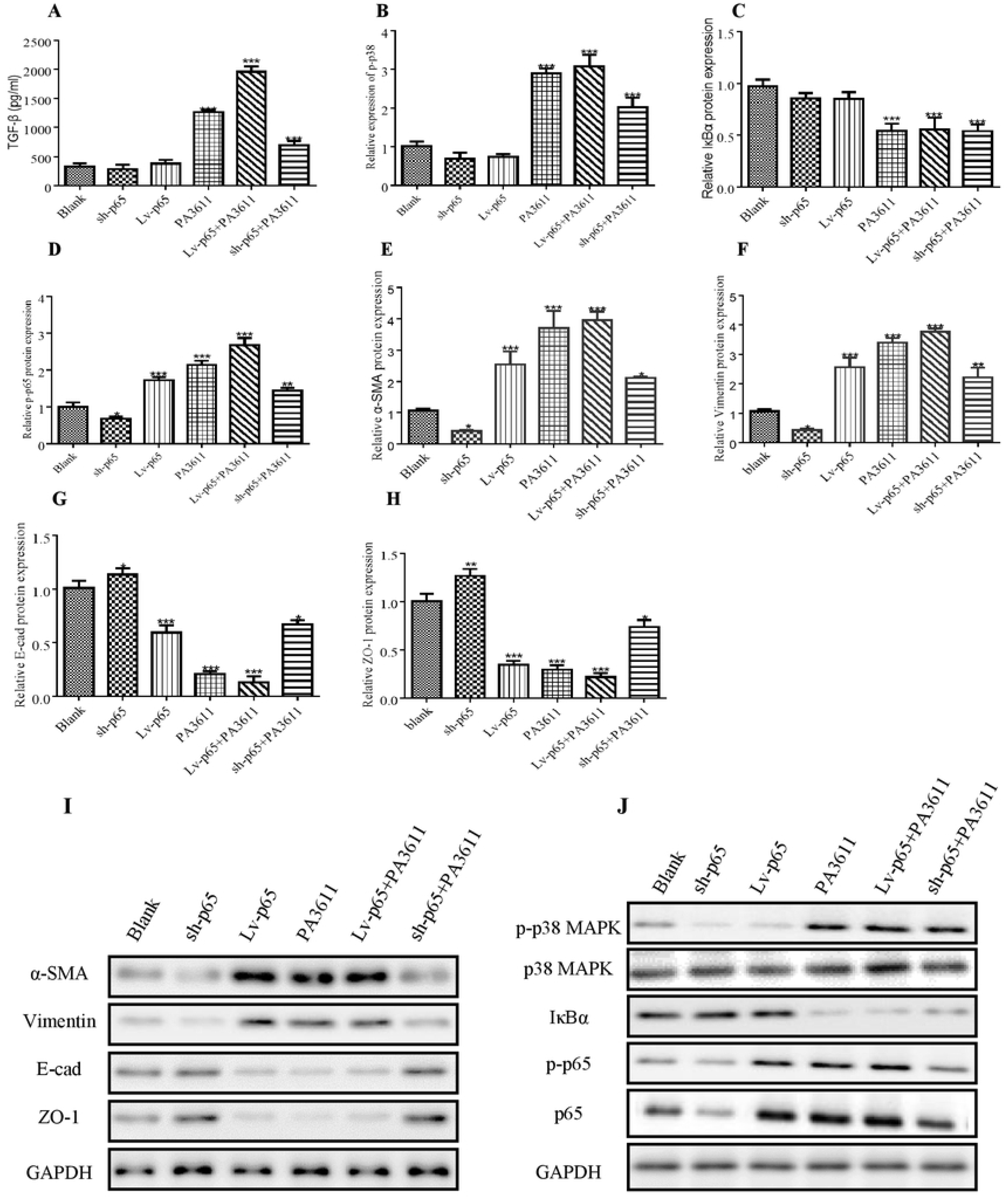
Effects of p65 overexpression or knockdown on the expression of TGF-β1, p38 phosphorylation, IκB and EMT relative markers. 16HBE cells were transfected with a control vector (indicated with Blank), a p65 overexpression vector (indicated with p65), or a specific siRNA directed against p65 (indicated with sh-p65). After 24h post transfection, cells were infected with PBS (negative control) or PA3611(30μg/ml) for another 24 hs, then cells culture supernatants were collected for TGF-β1 levels evaluated by enzyme-linked immunosorbent assay (Fig. 3A) and cells were collected for the protein expression of p38, IκBα, p65 and EMT relative markers (Fig. 3B-J). The results were representative of three independent experiments. Data were presented as the means±SD. *, P<0.05, **, P<0.01, ***, P<0.001, vs. the control vector or the nonspecific siRNA interference group of the same treatment.

### miR-3065-5p and miR-6802-3p downregulate the expression of IκBα and upregulate p65 phosphorylation without affecting TGF-β1expression and p38 phosphorylation

As miR-3065-5p and miR-6802-3p were observed to be upregulated in PA3611 infected bronchial epithelial cells, we hypothesized that they participated in regulating of NF-κB activation or p38 phosphorylation. Therefore, 16HBE cells were transfected with miR-3065-5p or miR-6802-3p mimics or with miR-3065-5p or miR-6802-3p inhibitors to evaluate their potential regulatory activities (Suppl Fig. 3–6). The results showed that both miR-3065-5p and miR-6802-3p downregulated the expression of IκBα and upregulate p65 phosphorylation, and this effect was enhanced by their combined activity or plus with PA3611(Fig. 4C-E). When miR-3065-5p and miR-6802-3p were knocked down, the level of IκBα was upregulated and p65 phosphorylation was decreased, however, pulsed with PA3611, the expression of the above indicators reversed (Fig. 5C-E). Both miR-3065-5p and miR-6802-3p upregulated the expression of α-SMA and Vimentin, and decreased the levels of E-cadherin and Zo-1from gene and protein levels, moreover, the expression levels changed more significantly after treated with PA3611 plus miR-3065-5p and miR-6802-3p (Suppl Fig. 3F-I, Suppl Fig. 4A-E, Suppl Fig. 5F-I, Suppl Fig. 6A-E). Neither miR-3065-5p nor miR-6802-3p had an effect on TGF-β1expression and p38 phosphorylation, but when added PA3611, TGF-β1 and p38 both increased (Fig. 4A-B, Fig. 5A-B). This finding suggested that miR-3065-5p and miR-6802-3p function as negative regulators upstream of IκBα but act downstream of p38 or had no obvious relationship with p38. To verify the hypothesis, the relationship between miRNAs (miR-3065-5p and miR-6802-3p) and IκBα were analyzed using Miranda. The IκBα-UTR sequence was observed to contain a binding site for miR-3065-5p and miR-6802-3p (Fig. 4F-G). A dual-luciferase reporter assay was subsequently performed to verify the interaction between miRNAs (miR-3065-5p and miR-6802-3p) and IκBα. The results showed that both miR-3065-5p and miR-6802-3p suppress luciferase expression, once the putative binding sites were mutated, miR-3065-5p or miR-6802-3p failed to significantly downregulate the expression of luciferase (Fig. 4H-I). These findings indicated that miR-3065-5p and miR-6802-3p binding sites were present in the IκBα-UTR, and that miR-3065-5p and miR-6802-3p directly downregulated the expression of IκBα. The binding was specific, because the luciferase expression was not affected by the control miRNA.

**Figure 4:**
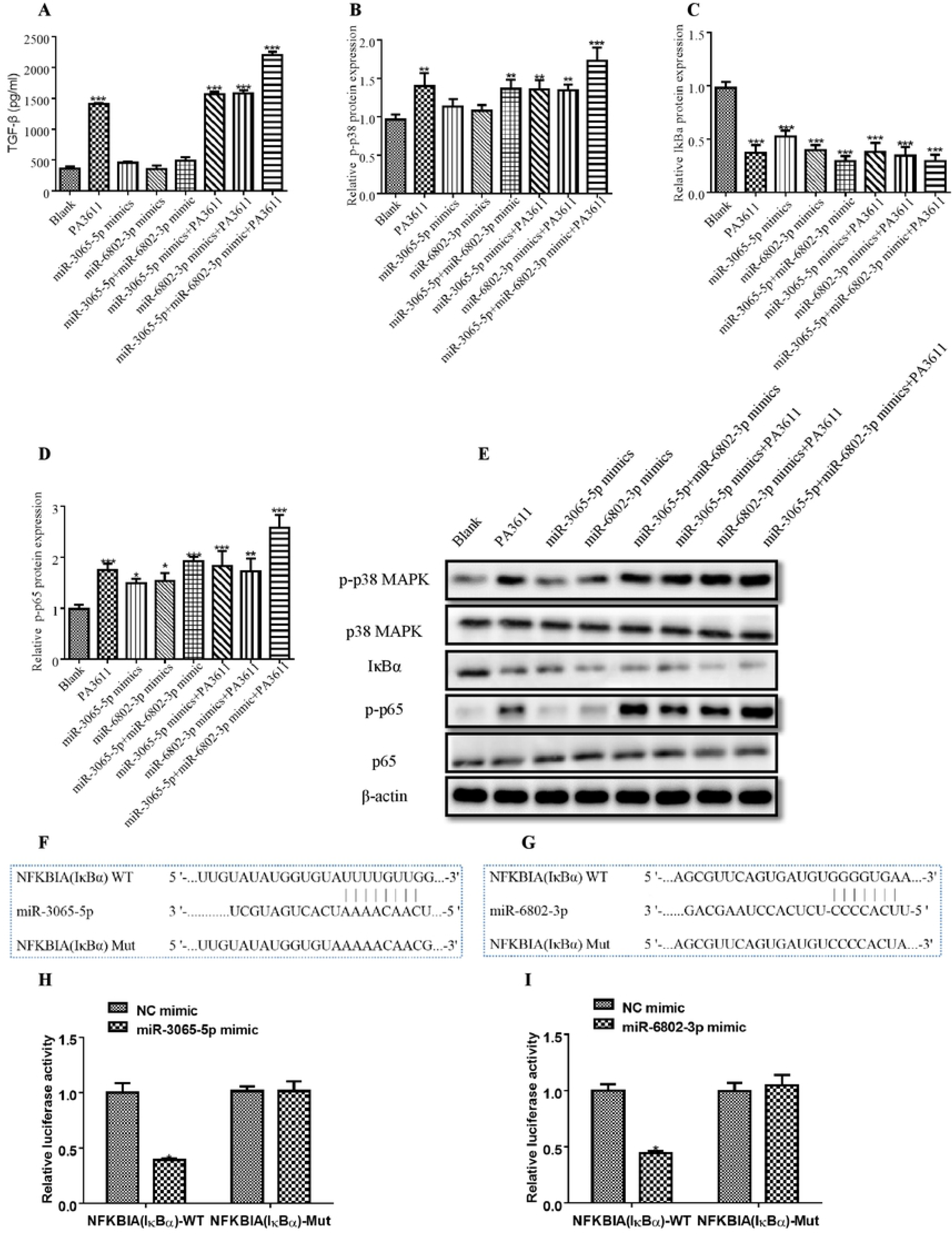
Effect of miR-3065-5p and miR-6802-3p intervention on the expression of TGF-β1, p38, IκBα and p65. 116HBE cells were transfected with a negative control miRNA mimic (indicated with Blank), miR-3065-5p, miR-6802-3p mimics, or both miR-3065-5p and miR-6802-3p mimics. After 24h post transfection, cells were infected with PBS (negative control) or PA3611(30μg/ml) for another 24 hs, then cells culture supernatants were collected for TGF-β1 levels evaluated by enzyme-linked immunosorbent assay (Fig. 4A) and cells were collected for the protein expression of p38, IκBα and p65(Fig. 4B-E). Sequence alignment of miR-3065-5p and miR-6802-3p and their conserved target sites in the IκBα-UTR were shown (Fig. 4F-G). Luciferase activity was measured in 16HBE cells with a dual-luciferase reporter assay. The cells were cotransfected with a plasmid expressing miR-3065-5p or miR-6802-3p mimic or a control miRNA (indicated with NC) and a vector expressing IκBα-UTR WT or IκBα-UTR MUT. Firefly luciferase activity was normalized to Renilla luciferase activity (Fig. 4H-I). The results were representative of three independent experiments. Data were presented as the means±SD. *, P<0.05, **, P<0.01, ***, P<0.001, vs. the control vector or the nonspecific siRNA interference group of the same treatment.

**Figure 5:**
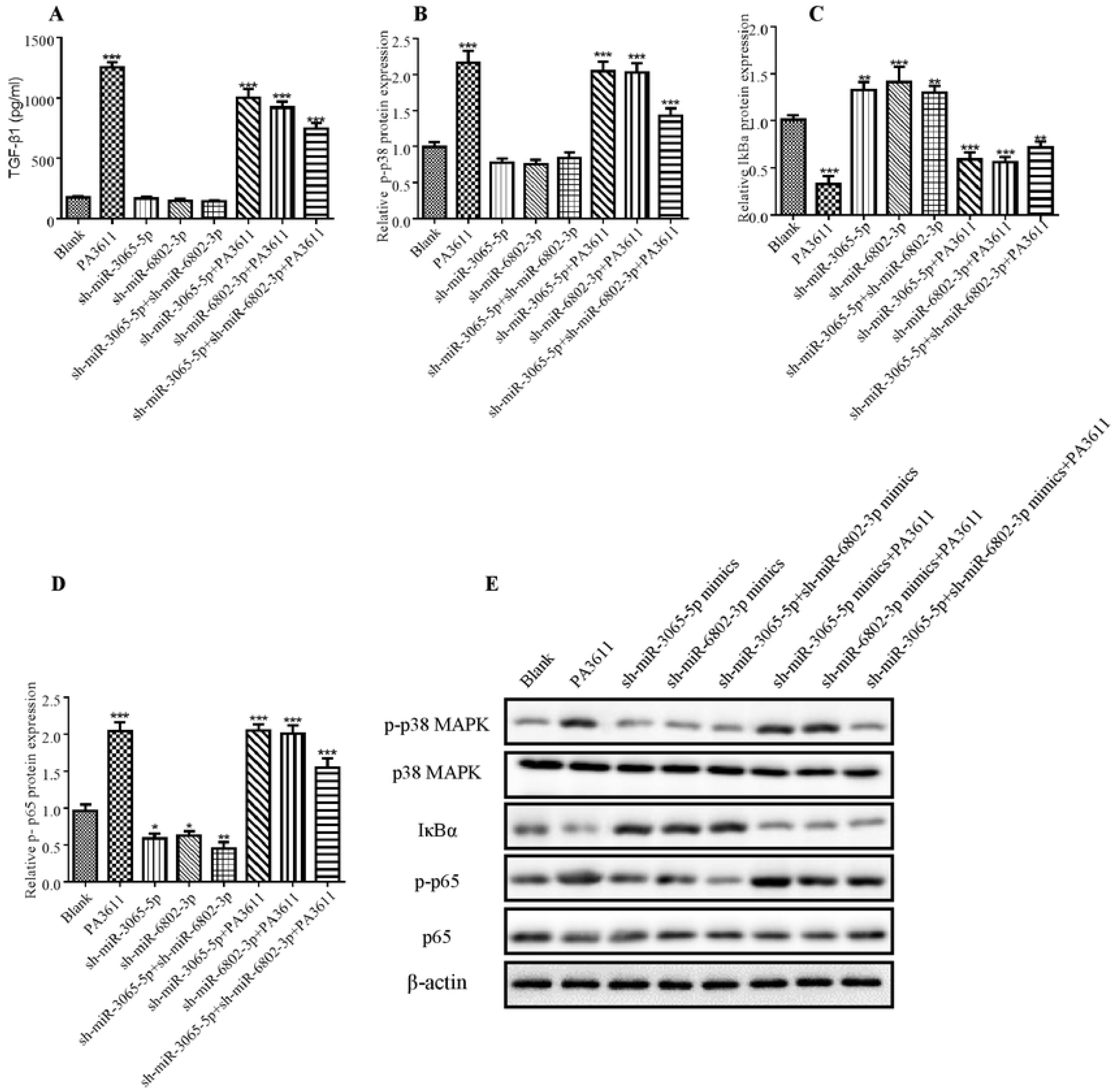
Effect of miR-3065-5p and miR-6802-3p knockdown on the expression of TGF-β1, p38, IκBα and p65. 16HBE cells were transfected with a negative control miRNA inhibitor (indicated with Blank), a miR-3065-5p inhibitor (indicated with sh-miR-3065-5p), a miR-6802-3p inhibitor (indicated with sh-miR-6802-3p), or both the miR-3065-5p and miR-6802-3p inhibitors (sh-miR-miR-3065-5p +sh-miR-6802-3p). After 24h post transfection, cells were infected with PBS (negative control) or PA3611(30μg/ml) for another 24 hs, then cells culture supernatants were collected for TGF-β1 levels evaluated by enzyme-linked immunosorbent assay (Fig. 5A) and cells were collected for the protein expression of p38, IκBα and p65(Fig. 5B-E). The results were representative of three independent experiments. Data were presented as the means±SD. *, P<0.05, **, P<0.01, ***, P<0.001, vs. the control vector or the nonspecific siRNA interference group of the same treatment.

### p38 regulates miRNAs expression, IκBα and p65 levels and EMT markers expression without affecting TGF-β1

In order to identify the function of p38 in PA3611 induced bronchial epithelial cells epithelial-mesenchymal transition, 16HBE cells were transfected with pcDNA3.1/p38 cDNA or a specific siRNA directed against p38. The results showed that overexpression of p38 resulted in upregulation of miR-3065-5p, miR-6802-3p and p65 phosphorylation, and decreased the level of IκBα, meanwhile, p38 overexpression upregulated the expression of α-SMA and Vimentin, and decreased the levels of E-cadherin and Zo-1from gene and protein levels (Fig. 6C-J, Suppl Fig. 7A-I), moreover, the expression levels changed more significantly after treated with PA3611 plus p38 overexpression (Fig. 6C-J, Suppl Fig. 7A-I). In addition, knockdown of p38 caused the downregulation of miR-3065-5p, miR-6802-3p and p65 phosphorylation, and increased that of IκBα (Fig. 6C-J, Suppl Fig. 7A-I). However, neither overexpression nor knockdown of p38 had not affect the secretion of TGF-β, in contrast, TGF-β expression significantly increased after treated with PA3611 plus overexpression or knockdown of p38 (Fig. 6A). These data indicated that PA3611 promotes bronchial epithelial cells epithelial-mesenchymal transition through TGF-β1 inducing p38/miRNA/NF-κB pathway.

**Figure 6:**
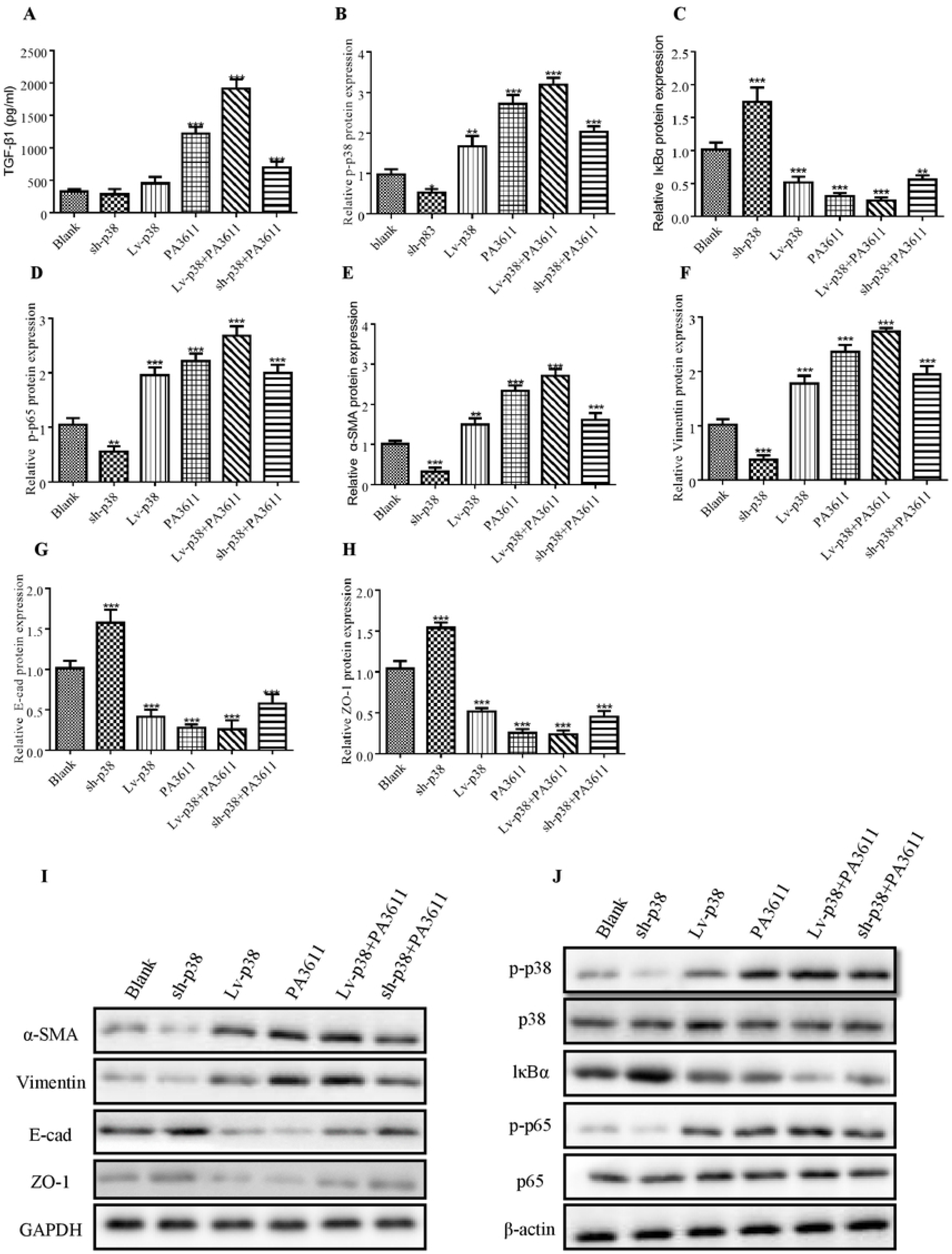
Effects of p38 overexpression or knockdown on the expression of TGF-β1, p38 phosphorylation, IκB, p65 and EMT relative markers. 16HBE cells were transfected with a control vector (indicated with Blank), a p38 overexpression vector (indicated with p38), or a specific siRNA directed against p38 (indicated with sh-p38). After 24h post transfection, cells were infected with PBS (negative control) or PA3611(30μg/ml) for another 24 hs, then cells culture supernatants were collected for TGF-β1 levels evaluated by enzyme-linked immunosorbent assay (Fig. 6A) and cells were collected for the protein expression of p38, IκBα, p65 and EMT relative markers (Fig. 6B-J). The results were representative of three independent experiments. Data were presented as the means±SD. *, P<0.05, **, P<0.01, ***, P<0.001, vs. the control vector or the nonspecific siRNA interference group of the same treatment.

**Figure 7:**
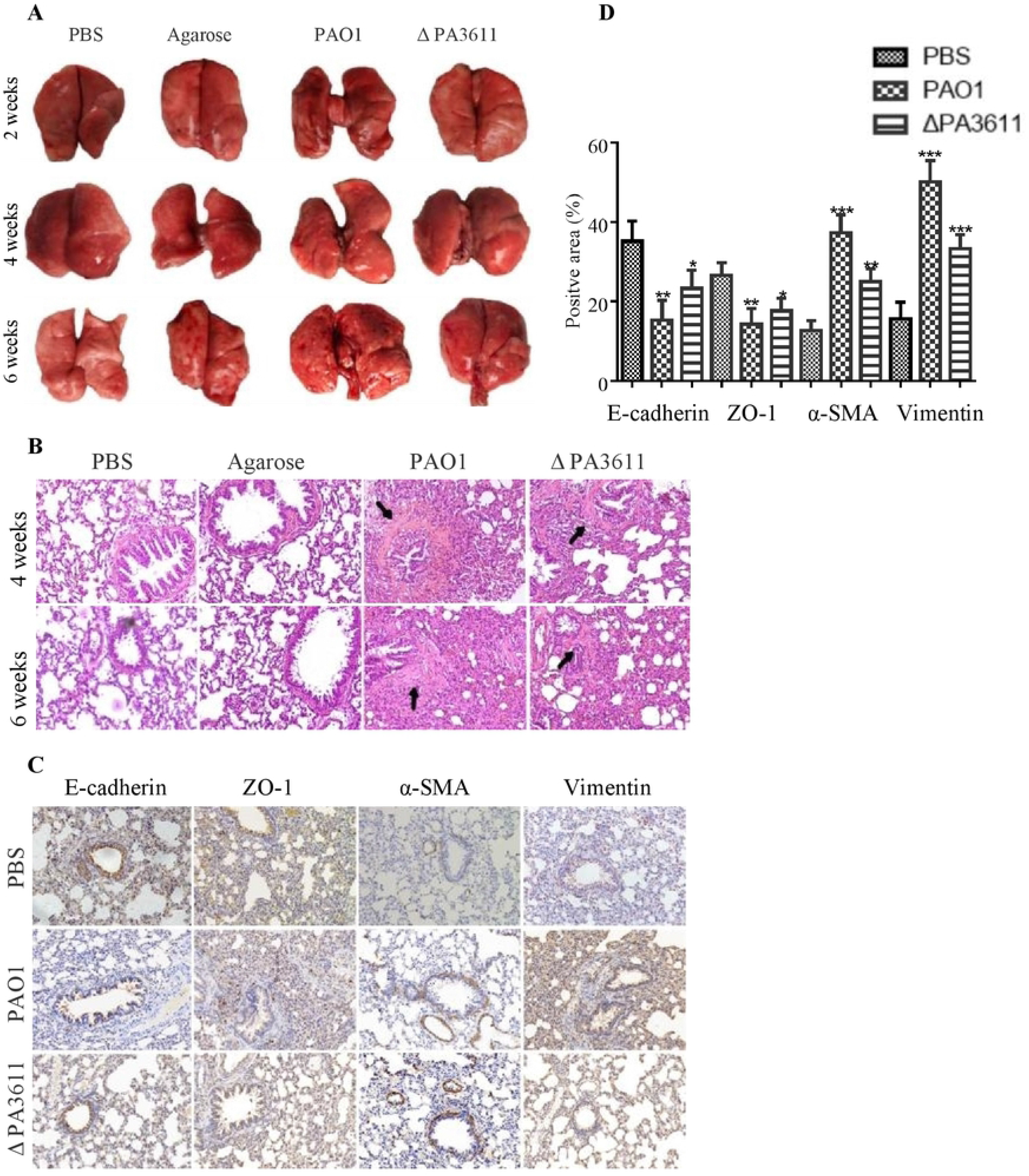
Effects of PA3611 on lung infection and epithelial mesenchymal transition (EMT) in vivo. 6-8 weeks old male Wistar rat were intratracheally infected with PAO1 or ΔPA3611bacterial strain. (The strain was coated with Agarose, Agarose and PBS as for control), lungs were collected at 2 weeks, 4 weeks or 6 weeks post infection. Overall gross view of lung tissues from four groups at 2 weeks, 4 weeks and 6 weeks post infection (A). HE staining (B) and Immunohistochemical staining of E-cadherin, ZO-1, α-SMA and Vimentin (C) Scale bar represents 100μm. Morphometric analysis of EMT relative proteins positive area were performed on immunohistochemical staining sections of lung tissues (D). The lung injury results are representative of three independent experiments. Data are presented as the means±SD, *, P <0.05, **, P <0.01, ***, P <0.001, vs. control group.

**Figure 8:**
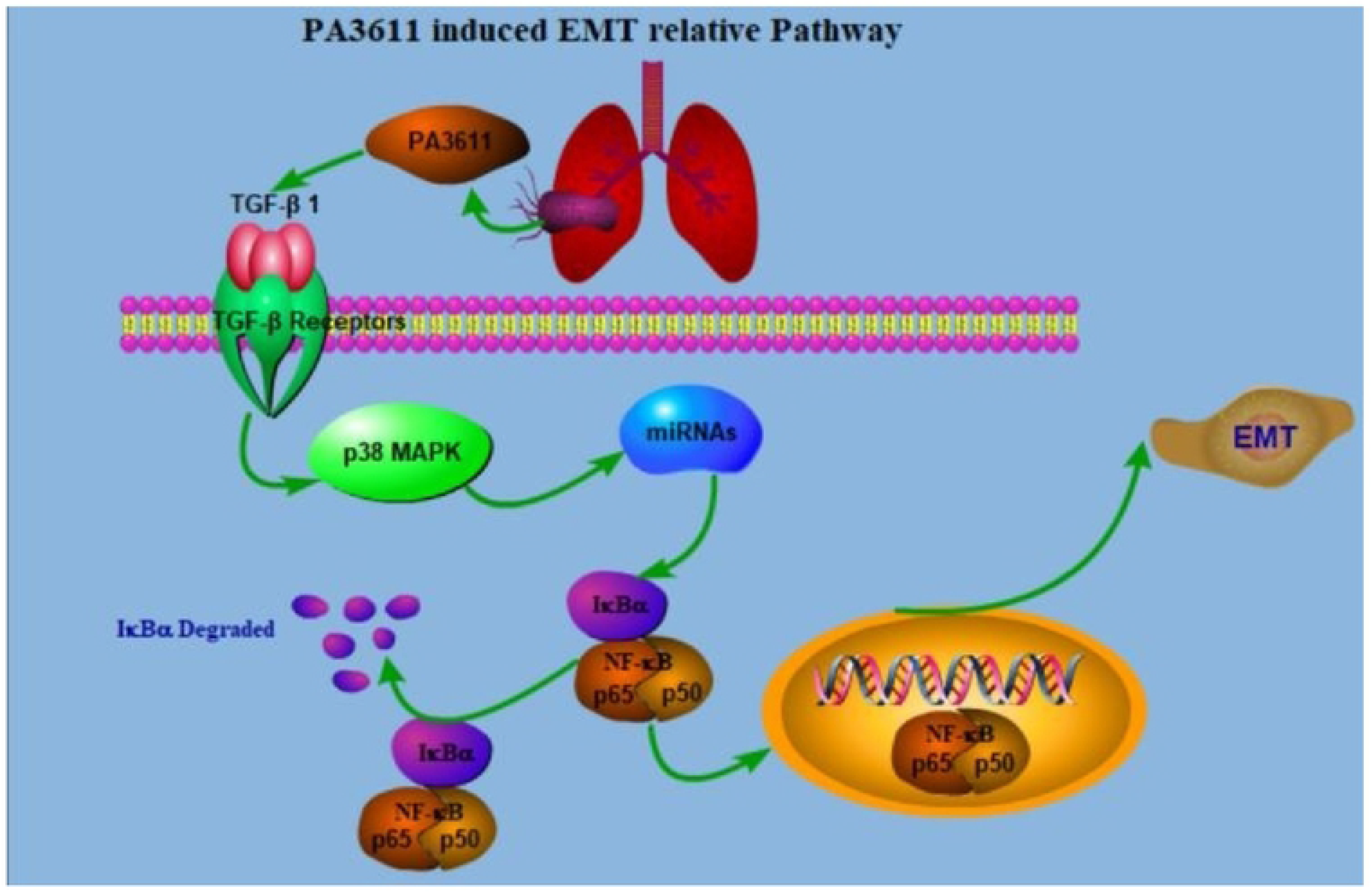
Schematic diagram of PA3611 induced EMT relative pathway.

### PA3611 promotes bronchial epithelial cells epithelial-mesenchymal transition in vivo

To verify the function of PA3611 in PA infection and epithelial-mesenchymal transition in vivo, 6-8 weeks old male rats were intratracheally infected with PBS, agarose, agarose coated PAO1 or agarose coated ΔPA3611. Lung morphology results showed mild hyperemia were observed locally in agarose coated PAO1 group 2 weeks post infection, at 4 weeks post infection, scattered nodules on the lung surface with local bleeding points were seen in agarose coated PAO1 or agarose coated ΔPA3611group, and these phenomena were more pronounced at 6 weeks post infection (Fig 6 A). HE staining revealed alveolar cavity collapse, intra-alveolar hemorrhage, partial pulmonary septum rupture, alveolar septum widening, bronchial lumen stenosis and deformation, smooth muscle proliferation and fibrosis around the trachea, and inflammatory cells infiltration in the lung interstitium from agarose coated PAO1 and agarose coated ΔPA3611 groups started from 4 weeks post infection, and more obviously at 6 weeks post infection (Fig 6 B). Epithelial-mesenchymal transition immunohistochemistry analyses confirmed that rats treated with agarose coated PAO1 and agarose coated ΔPA3611 both increased α-SMA and Vimentin proteins expression and deceased the expression of E-cadherin and ZO-1(Fig 6 C, D), these were in accordance with the in vitro results and indicating that PA3611 has a role on causing lung infection and promoting bronchial epithelial cells epithelial-mesenchymal transition.

## Discussion

PA is one of the main pathogens of chronic airway structural diseases such as cystic fibrosis, COPD, bronchiectasis, and bronchiolitis obliterans[23]. Among these patients with chronic lung diseases, COPD has a huge consumption of medical resources, and its cause of death ranks fourth in the world, and is expected to become the third leading cause of death by 2020[24,25]. Studies have found that the colonization of PA in the airway can cause chronic airway damage in COPD patients, leading to a significant declined in the forced expiratory volume in one second (FEV1) of COPD patients, suggesting that PA infection can promote COPD, bronchiectasis and cystic fibrosis patients with airway wall thickening, luminal narrowing and other fibrotic symptoms, which aggravate the irreversible airflow limitation of the patients’ airway[26,27,28,29]. Although the specific mechanism of PA infection causing related airway remodeling is still unclear, it is clear that EMT plays a key role in the process of bronchial fibrosis[30]. In this study, we demonstrated that PA3611 could play a crucial role in PA-associated airway remodeling by stimulating TGF-β production.

The virulence factors produced by PA have the functions of maintaining the growth and metabolism of PA itself, promoting the colonization and growth of bacteria in the host, and helping bacteria to adhere to infection or destroy host cells. PA3611 is a virulence factor secreted under PA infection, and is expressed in multiple subgroups of PA. Studies have shown that the lysate of inactivated PA can induce EMT in A549 cells[31], however, so far no studies have been focused on bronchial epithelial cells EMT caused by virulence factors secreted by PA infection.

In this study, a recombinant protein PA3611 was constructed through prokaryotic expression, which has a tendency to induce EMT in bronchial epithelial cells. We first studied its effect on the proliferation of 16HBE and RTE cell lines, and results showed that PA3611 could inhibit cell proliferation, and the inhibition effect was time and dose dependent.

Transmission electron microscopy results showed that after PA3611 stimulated above cells for 72hs, in addition to the enlargement of intercellular space and the disappearance of intercellular connections, filamentous pseudopodia appeared, and the ultrastructural manifestation of nuclear abnormity, peri-nuclear space broadening, mitochondria swelling and distortion, and intracellular glycogen increasing, which was consistent with the previously reported changes in the ultrastructure of EMT[32,33,34].

EMT is a pathological process in which cells lose their original cell polarity and transform into mesenchymal cell morphology and characteristics. Original epithelial cell markers, such as E-cad and ZO-1 expressions decreased, and mesenchymal cell markers, such as α-SMA, N-cad and Vimentin expressions increased, and the activity of some matrix metalloproteinases (MMPs) also increased. In this study, by detecting the expression of 16HBE and RTE-related markers at the time points of PA3611 protein stimulation for 24, 48 and 72 hs, the expression of epithelial cell markers E-cad and Zo-1 decreased, while mesenchymal cell markers Vimentin and α-SMA were increased, which indicated that the cells had been undergone EMT changes.

TGF-β1 has been considered as a key inducer of EMT and has a central role in regulating fibrosis process through several different mechanisms[35,36,37]. Studies also have found that the nuclear transcription factor NF-κB can regulate the process of EMT by controlling the Smad-independent gene network[38], reducing the activation of NF-κB can alleviate alveolar epithelial cells EMT induced by bleomycin, and inhibiting NF-κB can prevent bronchial epithelial cells EMT occurrence induced by tumor necrosis factor ligand superfamily member 14 (TNFSF14)[39,40]. NF-κB binds with IκBα in a resting state to form an inactive complex, which makes it stay in the cytoplasm, when the cell is stimulated by certain external signals, the complex is decomposed, and NF-κB p65 is phosphorylated[41]. In this study, intracellular NF-κB p65 phosphorylation levels were significant higher than that in control group at the time of PA3611 stimulation for 24 hs, when added TGF-β1 neutralizing antibody, compared with PA3611 and TGF-β1 stimulated groups, the expression of E-cadherin and ZO-1 increased, while Vimentin and α-SMA decreased, and consistent with the expression levels in control group, meanwhile, and the phosphorylation level of NF-κB p65 was reduced. These results indicated that blocking TGF-β1 could inhibit the occurrence of PA3611-induced EMT, which suggesting that PA3611-induced EMT changes in epithelial cells were achieved by stimulating of TGF-β1 secretion.

It is well known that NF-κB is an important intersection for a variety of signaling pathways, and many factors play roles in NF-κB activation, in recent years, the activating effect of microRNA (miRNA) has attracted much attention[42,43]. miRNA is a kind of single-stranded small molecule RNA with a length of about 21~25 nucleotides that widely exists in eukaryotes. It does not encode proteins, but mature miRNA can specifically degrade or hinder the translation of target mRNA by binding to its target gene in the 3’untranslated region (3’UTR)[44]. Studies have shown that miR-424 can negatively regulate myofibroblast differentiation and achieve an inhibitory effect on EMT[45], and miR-135a can mediate EMT by targeting specific cytokines[46]. These findings indicate that miRNAs play a specific and important role in regulating the EMT in related cells. Feng et al. [47] found that miRNA-126 can promote the activation of NF-κB by targeting the 3’-untranslated region (3’-UTR) of IκBα, leading to hepatic stellate cell fibrosis, and Yao et al.[48] found that miRNA-891a-5p can directly target and inhibit IκBα and promote NF-κBp65 phosphorylation in relevant studies on abnormal angiogenesis. These studies suggest that miRNA s can promote NF-κB p65 activation by targeted inhibition of IκBα. In our study, the levels of miR-3065-5p and miR-6802-3p were significantly higher in PA3611 treated with 16HBE cells compared with untreated cells. Further investigation suggested that 3065-5p and miR-6802-3p inhibited the expression of IκBα and the binding sites are present in the IκBα-UTR, indicating that 3065-5p and miR-6802-3p are negative regulators of IκBα.

According to our results, we conclude that PA bacteria infect bronchial epithelial cells by secreting PA3611 protein which activates p38 MAPK signaling molecules through the secretion of TGF-β1, then stimulating the up-expression of 3065-5p and miR-6802-3p, the later targeting on the IκBα gene and inhibiting its expression and promoting the activation of NF-κB p65, finally induce EMT occur in bronchial epithelial cells. Therefore, in our study, we elucidate PA3611 can promote bronchial epithelial cells epithelial-mesenchymal transition through TGF-β1 inducing p38/miRNA/NF-κB pathway, and expect to provide a new potential target for the clinical treatment of bronchial EMT and fibrosis caused by chronic PA infection.

## Materials and methods

### Recombinant PA3611 protein *in vitro*

A DNA fragment encoding PA3611 was obtained from Qiangyao Biological Company (Suzhou, China) and the oligonucleotide primers were designed with Primer 5.0 (Premier, Canada) (Table S1). The DNA template and primers were synthesized by Qiangyao and PCR was performed with the template described above. The condition of PCR amplification used was as follows: 94°C for 5min, followed by 30 cycles of 96°C for 25s, 58°C for 25s, and 72°C for 1min. The PCR product was cut and purified via DNA gel extraction kit (AXYGEN, NY, USA). The purified PCR product and pET-28a(+) (Novagen, Germany) were digested with BamH I and Xho I (NEB) (Thermo Scientific, DE, USA) and ligated together with T4 DNA ligase (Thermo Scientific). The pET-28a(+) vector containing the PA3611 encoding gene was transformed into *E. coli* strain BL21(DE3) (Novagen) and subsequently grown on Luria-Bertani (LB) agar plates containing 50μg/ml kanamycin at 37°C overnight. Positive clones were selected for enzyme digestion and sequencing identification. The sequence-corrected plasmids were transformed into *E. coli* BL21(DE3) and plated onto solid medium, containing kanamycin, and incubated overnight at 37°C at 250r/min. 1% of overnight bacteria were transferred to LB medium containing kanamycin at 37°C at 250 r/min for 3 h, and then add 0.5 mmol/L IPTG at 20°C for 12 h. Next, the culture was centrifuged and cells were resuspended and lysed by sonication. The sonicated sample was centrifuged and the supernatant of the cell lysate was applied to a Ni-IDA resin (Qiangyao). The protein was subsequently eluted and collected for SDS-polyacrylamide gel electrophoresis analysis. A protein with the predicted mass of PA3611 was concentrated through ultrafiltration (molecular weight cutoff 16kDa). The concentration of the protein was determined by the Bradford method. Subsequently, a western blotting was performed to examine the purified recombinant PA3611 protein via the N-His Tag (His Tag antibody; Qiangyao). then filtered and sterilized with a 0.22μm sterile membrane, and the protein was stored at −80°C.

### Cells and Bacterial culture

Human bronchial epithelial ceAll line 16HBE and rat bronchial epithelial cell line RTE were both preserved in our laboratory. Cells were cultured in RPMI-1640 medium (Gibco BRL, Grand Island, NY, USA) containing 10% fetal bovine serum (FBS, Gibco BRL), penicillin (100U/ml), streptomycin (100μg/ml), and L-glutamine (2mM), and placed in cell incubator with constant temperature of 37°C and 5% CO_2_ under a humidified atmosphere, the medium was changed every other day, and passage was carried out in two to three days according to cells growth situation, logarithmic growth phase cells were used for experiments.

Pseudomonas aeruginosa standard strain PAO1 (ATCC15692) was kindly donated by Professor Jinfu Xu from Shanghai Chest Hospital. Pseudomonas aeruginosa knockout strain ΔPA3611 was purchased from Guangzhou Nuojing Biotechnology Company, both strains were cultivated on Cetrimide-agar (Merck, Darmstadt, Germany) plates. For the experiments single colonies of bacteria were inoculated in Luria-Bertani broth (Merck) and incubated over night at 37°C and 140 rpm and stored at −80°C. Before infecting animals, stock solutions of PA were thawed, washed, and diluted in sterile distilled water to a specific concentration.

### Cells transfection test

The fragments encoding the P65 and P38 alleles genes were obtained from GenScript and cloned into pcDNA3.1 (Invitrogen, Carlsbad, CA, USA). 16HBE and RTE cells were transiently transfected with pcDNA3.1/p65 cDNA, pcDNA3.1/p38 cDNA, control pcDNA3.1, miR-155 mimic (GenePharma, Shanghai, China), miR-99b mimic (GenePharma), or a negative control (NC) miRNA (GenePharma) using Lipofectamine 3000 (Invitrogen) according to the manufacturer’s instructions. Cells were incubated for 24h at 37°C before being used for further analysis.

siRNAs targeting NF-κB p65 subunit mRNA or p38 mRNA, a random non-coding siRNA, a miR-3605-5p inhibitor, a miR-6082-3p inhibitor and an NC mRNA were synthesized by Gene Pharma. 16HBE cells were transfected with the above siRNAs or miRNAs using Lipofectamine 3000 according to the manufacturer’s instructions. A non-coding siRNA or an NC miRNA was used as a NC. The cells were transfected with above RNAs for 24 hs, then infected with PBS (negative control) or PA3611(30μg/ml) for another 24 hs and carried out further experiments. The corresponding sequences of these RNAs are shown in Table S2.

### Proliferation assay

16HBE and RTE cells were seeded in a 96-well cell culture plate at 6000 cells per well and cultured overnight, then added PA3611 with a protein concentration of 0 μg/ml, 5 μg/ml, 15 μg/ml, 30 μg/ml, 45 μg/ml, and use 10% FBS RPMI-1640 medium to adjust the total volume of each well to 100 μl (changed the culture solution every 24h and made the PA3611 protein concentration consistent with the first time). After incubating for 24, 48 or 72hs, the proliferation assay was performed using a CCK-8 kit (Dojindo Laboratories, Kumamoto, Japan) according to the manufacturer’s protocol.

### ELISA assay

In order to determine the secretion of active TGF-β1, cell-free supernatants were collected and used to evaluate the concentrations of TGF-β1 with human and rat TGF-β1 ELISA kit (R&D Systems, Minneapolis, MN, USA) according to the manufacturer’s instructions.

### Western blotting analyses

16HBE and RTE Cells incubating for different time-points were collected and lysed. The lysates were centrifuged, denatured, applied to an SDS polyacrylamide gel for electrophoresis and transferred to a polyvinylidene fluoride membrane. Then the membranes were blocked with 5% skimmed milk at room temperature for 1h, and immunoblotting was carried out using antibodies to NF-κB p65 (Abcam, MA, USA), NF-κB p-p65 (phospho S536, Abcam), p38 (Abcam), p-p38 (phospho Y182, Abcam), α-SMA (Abcam), Vimentin (Abcam), E-cadherin (Abcam), ZO-1 (Abcam) and glyceraldehyde-phosphate dehydrogenase (GAPDH, Abcam). After washing, the membranes were incubated with the appropriate horseradish peroxidase-conjugated secondary antibody at room temperature for 1h. The bands were visualized via chemiluminescence using an ECL kit (Thermo scientific) and photographed with a Tanon Multi-Imager. The density of the immunoreactive bands was measured using ImageJ (Scion Corporation, Frederick, MD, USA).

### Quantitative reverse transcriptase-PCR

Total RNA was extracted from the cells using a Total RNA Extraction Kit (Generay Biotech, Shanghai, China), then RNA was reverse-transcribed into cDNA using a RevertAid First Strand cDNA synthesis Kit (Thermo Scientific Fisher, Waltham, MA, UK) and the thermocycling program used was as follows: 37°C for 60 mins and 85°C for 5mins. Amplifications were performed in an iCycler using iQ SYBR Green supermix (Bio-Rad). Gapdh was amplified on the same plates and used to normalize the data. Each sample was prepared in triplicate and each experiment was repeated at least three times. The relative abundance of each gene was quantified using the 2^-ΔΔCt^ method. The PCR primers used are listed in Table S1. Total RNA for miR-3605-5p, miR-6082-3p, and U6 detection was extracted with a Total RNA Extraction Kit. Reverse transcription and PCR was performed using a BulgeLoopTM miRNA qRT-PCR Starter Kit, a Bulge-Loop™ miRNA qRT-PCR Primer Set, and a U6 snRNA qPCR Primer Set (RiboBio, Guangzhou, China) according to the manufacturer’s instructions. miRNA expression was quantified using the 2^-ΔΔCt^ method and U6 was used as an internal control.

### Dual-luciferase reporter assay

The wild-type (WT) and mutant (MUT) 3’-untranslated region (3’-UTR) fragment of IκBα **(human/rat)** was inserted into the pGL3-basic vector (firefly luciferase; Promega, Madison, WI, USA), which was obtained from General Biosystems (Anhui, China). 16HBE or RTE cells were co-transfected with IκBα-UTR-WT or IκBα-UTR-MUT plasmids along with miR-3605-5p or miR-6082-3p, mimics or scramble oligonucleotides using Lipofectamine 3000. A reporter vector carrying the WT or MUT sequences of IκBα-UTR was assayed for luciferase expression using the Dual-Luciferase^®^ Reporter Assay System (Promega) following the manufacturer’s instructions. For data analysis, firefly luciferase activity was normalized to the corresponding Renilla luciferase activity.

### Preparation of Agarose-coated bacteria

A 2% agarose solution (with PBS) of 100 ml was prepared. 3.0g of tryptic soy broth (TSB) was added with 100 ml distilled water, and autoclaved at 121 °C for 15 min. Pseudomonas aeruginosa strains (PAO1 and ΔPA3611) were inoculated into solid medium for overnight at 37°C. Then selected colonies were inoculated into 5 ml TSB and cultured overnight, 1 ml of the overnight culture was put into a flask containing 10 mL TSB, and incubated at 37°C and 250 RPM until the logarithmic phase, then the bacterial culture was added to 10 mL preheated (48°C) agarose solution. Mixed quickly by vortexing and poured the agarose bacteria solution into the preheated paraffin oil and stirred at 500 rpm for 5 minutes, then cooled and removed the excess paraffin oil with vacuum pump and stored at 4°C for use.

### Infection of rats *in vivo*

6 to 8 weeks old male Wistar rats (Cavens Lab Animal Company, Changzhou, China) were housed in a bio-safety level III animal facility under specific pathogen-free conditions. All animal experimental procedures were approved by the Institutional Animal Ethics Committee of the Second Affiliated Hospital of Nanjing Medical University (No.2014KY050) and were carried out in strict accordance with the Nanjing Medical University’s guidelines for the use of laboratory animals. All rats were divided into four groups: control group (PBS group), agarose group, agarose coated PAO1 group (PAO1 group) and agarose coated ΔPA3611 group (ΔPA3611 group). The lung tissues were collected for histological and immunohistochemical staining at week 2, week 4, and week 6 respectively after intratracheal injection of agarose-coated suspension or PBS 120ul every two weeks for a total of 3 infections.

### Histology and Immunohistochemical staining

Lung lobes were collected and fixed with 4% paraformaldehyde overnight, and embedded in paraffin. H&E-stained tissues were assessed via a pathology analysis. Lung injury was estimated by the percentage of the lesion area in the total lung area using an ImagePro macro. Immunohistochemical staining of α-SMA, Vimentin, E-cadherin, ZO-1were performed using tissue sections that were dewaxed and rehydrated. Antigen retrieval was performed using a proteinase K and hot citric acid buffer treatment as needed. The restored sections were incubated with primary antibodies overnight at 4°C, after rinsing with Tris-buffered saline for 15minutes, sections were incubated with secondary antibody (biotinylated goat anti-rabbit IgG, Sigma). Sections then were washed and incubated with the Vect astain Elite ABC reagent (Vector Laboratories, Burlington, Ontario, Canada) for 45minutes. Staining was developed using 3,3-diaminobenzidine (2.5mg/mL) followed by counterstaining with Mayer’s hematoxylin. Images were taken with a Leica Microsystems Ltd microscope and were analyzed using ImagePro Plus 6.

### Transmission electron microscopy (TEM) examination

The16HBE and RTE cells were rinsed with ice-cold 0.1 M PBS (pH of 7.4) and centrifuged at 500 × g for 5 min at room temperature, after which the clear supernatants were removed. Cell pellets were fixed with 2.5% glutaraldehyde at least 30 min at 4 °C. After fixation, the treated cells were thoroughly washed in PBS and then post-fixed with 1% OsO4 for 1 h at room temperature. Then the specimens were embedded in Epon for 12 h at 35°C. Finally, 50-70 nm sections were stained with uranyl acetate (30 mins) and lead citrate (10 mins) at room temperature. Images were observed using a JEM-2000EX transmission electron microscope at 60 kV.

### Statistical analysis

All the presented results were expressed as mean ± SD, and confirmed in three independent experiments. The Student’s t-test was used to compare two groups and multiple groups were analyzed by performing one-way ANOVA. Statistical analyses were performed using SPSS 20.0 (IBM SPSS, Armonk, NY, USA). P<0.05 was considered to be significantly different.

## Acknowledgements

Not applicable.

## Authors’ contributions

The authors’ contributions are as follows: L. S., S.C. and G. F. conceived and designed the study; L. S., S. L., and G. F. conducted the research; X. C., K. D., Y. C., J. Y., Z. S., and X.D., analyzed and interpreted the data; and L. S., J.W., and G.F. wrote the manuscript; L.S., S.C., and G. F. revised the manuscript. All authors read and approved the final version of the manuscript.

## Availability of data and materials

The data and materials were as the contents we submitted, the other data and materials will be made available upon request.

## Ethics approval and consent to participate

All experimental procedures were approved by the Institutional Animal Ethics Committee of the Second Affiliated Hospital of Nanjing Medical University (No.2014KY050) and were carried out in strict accordance with the Nanjing Medical University’s guidelines for the use of laboratory animals.

## Consent for publication

Not applicable

## Supplemental Figure legends

**Supplemental figure 1:**
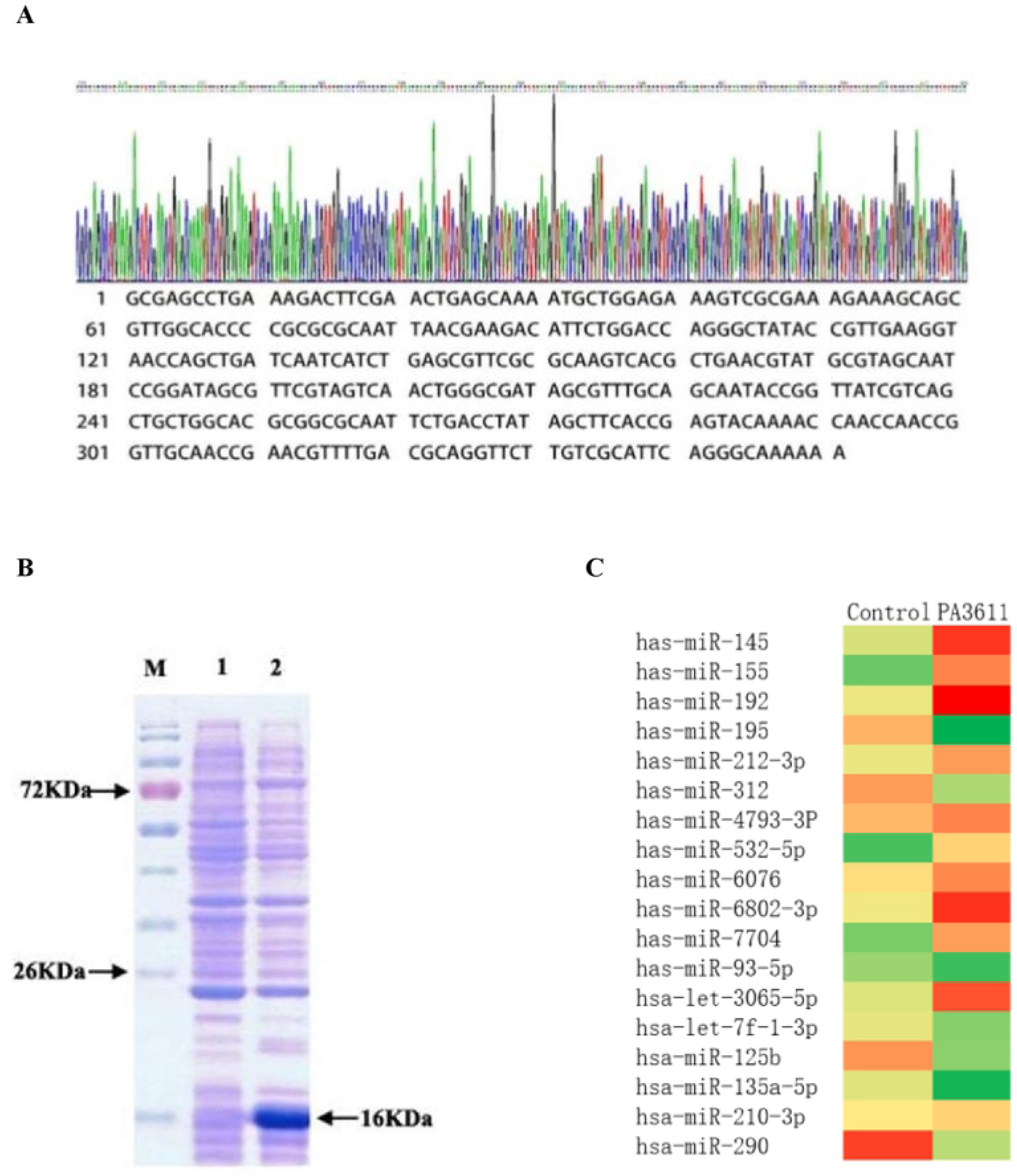
Heterologous expression and purification of recombinant PA3611 and miRNA array in PA3611 induced 16HBE cells. Sequencing results of recombinant plasmid PET-28a-PA3611 (Suppl Fig. 1A). Expression, purification and identification of PA3611 protein (M: Molecular weight marker of protein;1. Total protein of bacteria without induction;2: The total protein of the induced bacteria. Suppl Fig. 1B). miRNA array expression in PA3611 induced 16HBE cells (Suppl Fig. 1C).

**Supplemental figure 2:**
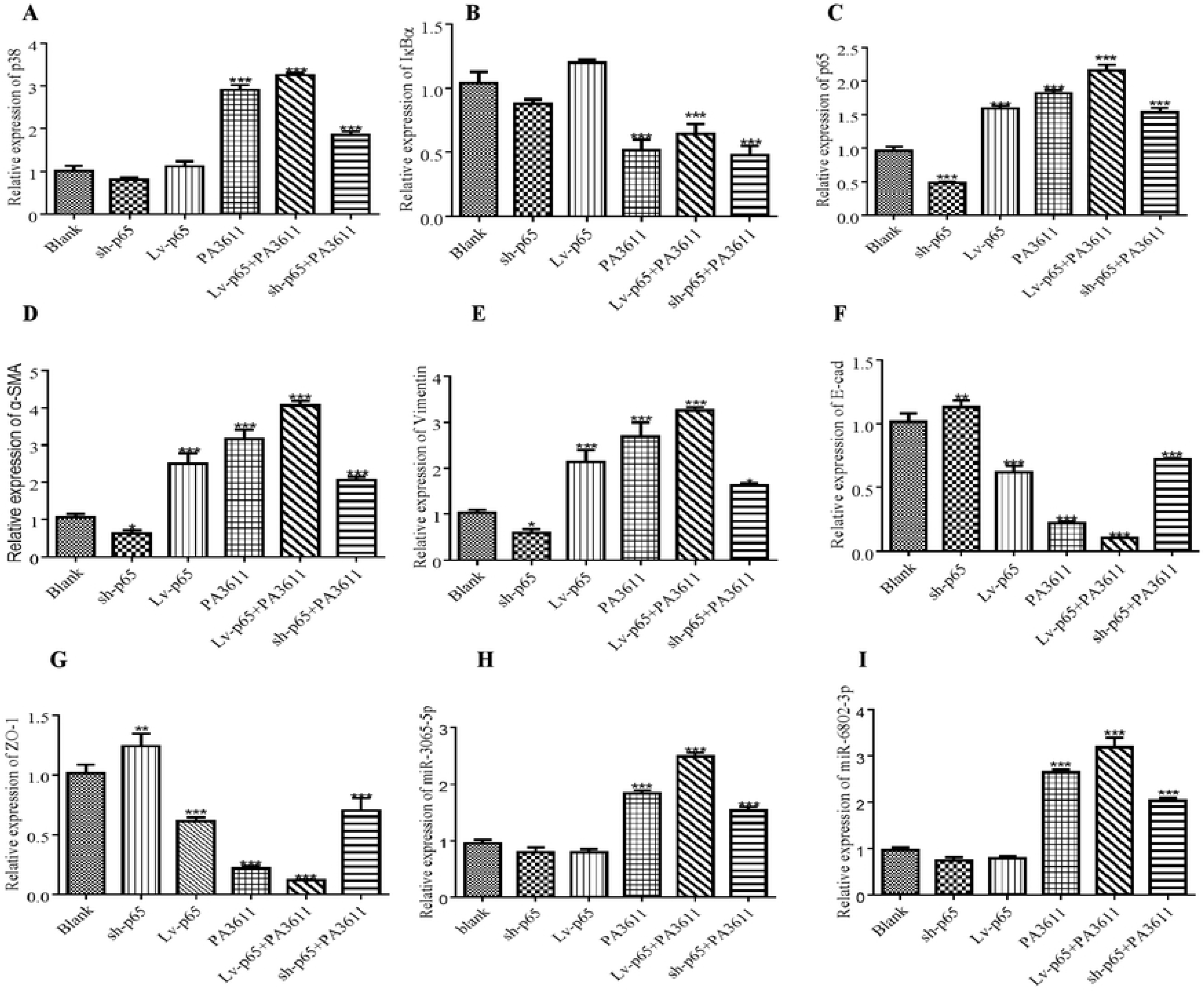
Effect of p65 intervention on the mRNAs expression of p38, IκBα, p65 and EMT relative markers. 16HBE cells were transfected with a control vector (indicated with Blank), a p65 overexpression vector (indicated with p65), or a specific siRNA directed against p65 (indicated with sh-p65). After 24h post transfection, cells were infected with PBS (negative control) or PA3611(30μg/ml) for another 24 hs, then cells were collected and detected for the mRNAs expression of p38, IκBα, p65 and EMT relative markers (Suppl Fig. 2A-G). The results were representative of three independent experiments. Data were presented as the means±SD. *, P<0.05, **, P<0.01, ***, P<0.001, vs. the control vector or the nonspecific siRNA interference group of the same treatment.

**Supplemental figure 3:**
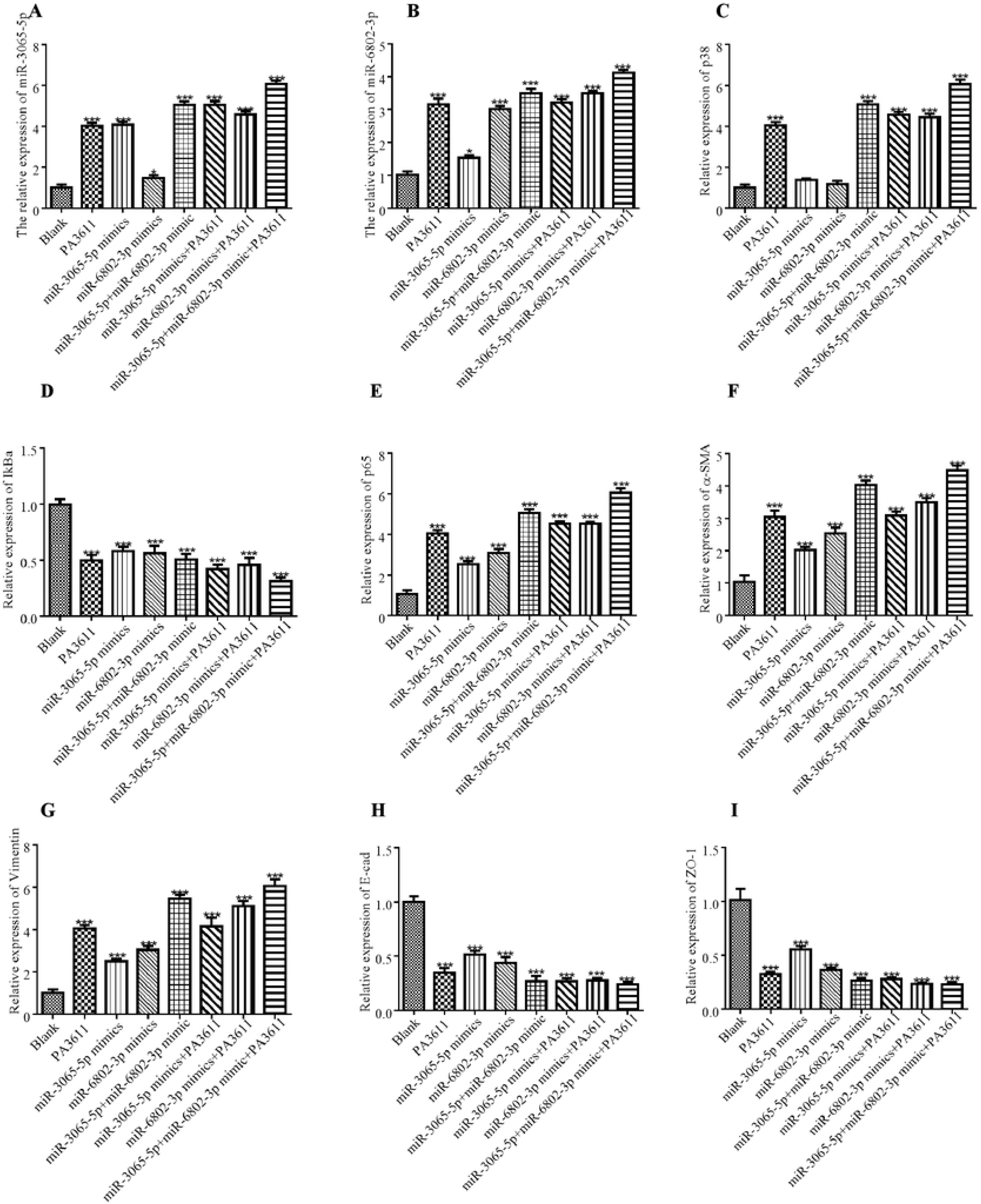
Effect of miR-3065-5p and miR-6802-3p intervention on the mRNAs expression of p38, IκBα, p65 and EMT relative markers. 16HBE cells were transfected with a negative control miRNA mimic (indicated with Blank), miR-3065-5p, miR-6802-3p mimics, or both miR-3065-5p and miR-6802-3p mimics. After 24h post transfection, cells were infected with PBS (negative control) or PA3611(30μg/ml) for another 24 hs, then cells were collected and detected for the mRNAs expression of p38, IκBα, p65 and EMT relative markers (Suppl Fig. 3A-I). The results were representative of three independent experiments. Data were presented as the means±SD. *, P<0.05, **, P<0.01, ***, P<0.001, vs. the control vector or the nonspecific siRNA interference group of the same treatment.

**Supplemental figure 4:**
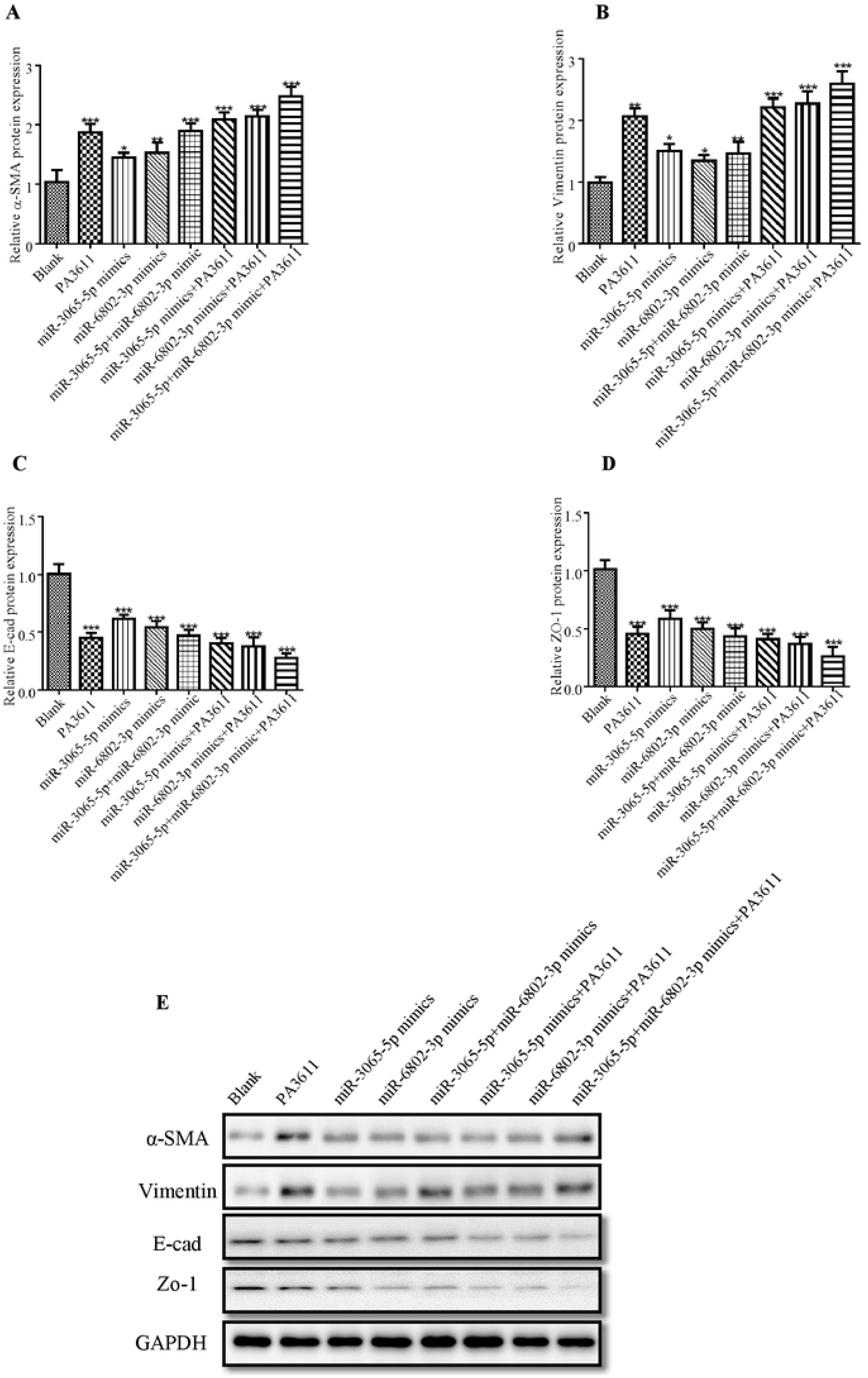
Effect of miR-3065-5p and miR-6802-3p intervention on the proteins expression of EMT relative markers. 16HBE cells were transfected with a negative control miRNA mimic (indicated with Blank), miR-3065-5p, miR-6802-3p mimics, or both miR-3065-5p and miR-6802-3p mimics. After 24h post transfection, cells were infected with PBS (negative control) or PA3611(30μg/ml) for another 24 hs, then cells were collected and detected for the proteins expression of EMT relative markers (Suppl Fig. 4A-E). The results were representative of three independent experiments. Data were presented as the means±SD. *, P<0.05, **, P<0.01, ***, P<0.001, vs. the control vector or the nonspecific siRNA interference group of the same treatment.

**Supplemental figure 5:**
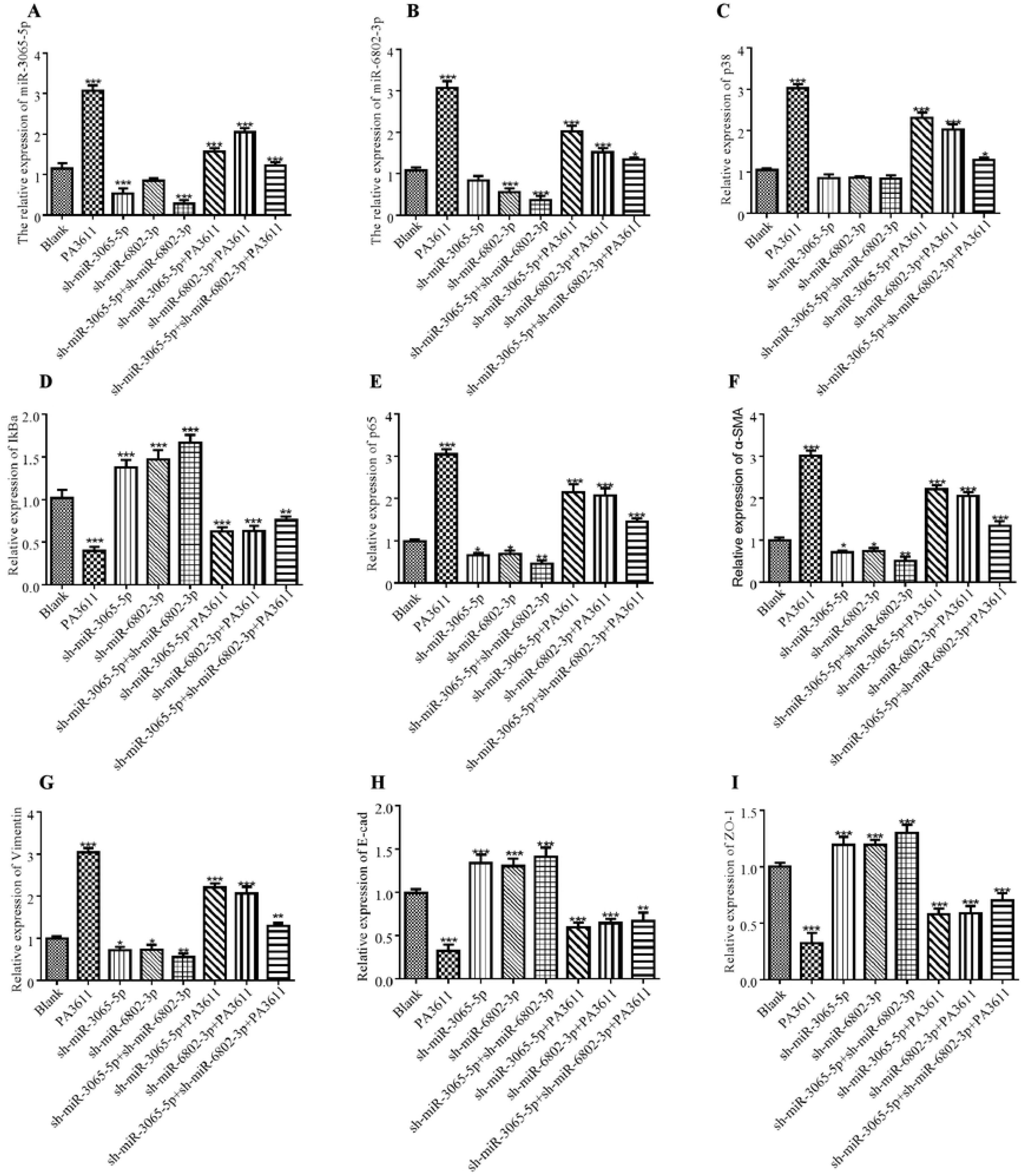
Effect of miR-3065-5p and miR-6802-3p knock down on the mRNAs expression of p38, IκBα, p65 and EMT relative markers. 16HBE cells were transfected with a negative control miRNA inhibitor (indicated with Blank), a miR-3065-5p inhibitor (indicated with sh-miR-3065-5p), a miR-6802-3p inhibitor (indicated with sh-miR-6802-3p), or both the miR-3065-5p and miR-6802-3p inhibitors (sh-miR-3065-5p+sh-miR-6802-3p). After 24h post transfection, cells were infected with PBS (negative control) or PA3611(30μg/ml) for another 24 hs, then cells were collected and detected for the mRNAs expression of p38, IκBα, p65 and EMT relative markers (Suppl Fig. 4A-I). The results were representative of three independent experiments. Data were presented as the means±SD. *, P<0.05, **, P<0.01, ***, P<0.001, vs. the control vector or the nonspecific siRNA interference group of the same treatment.

**Supplemental figure 6:**
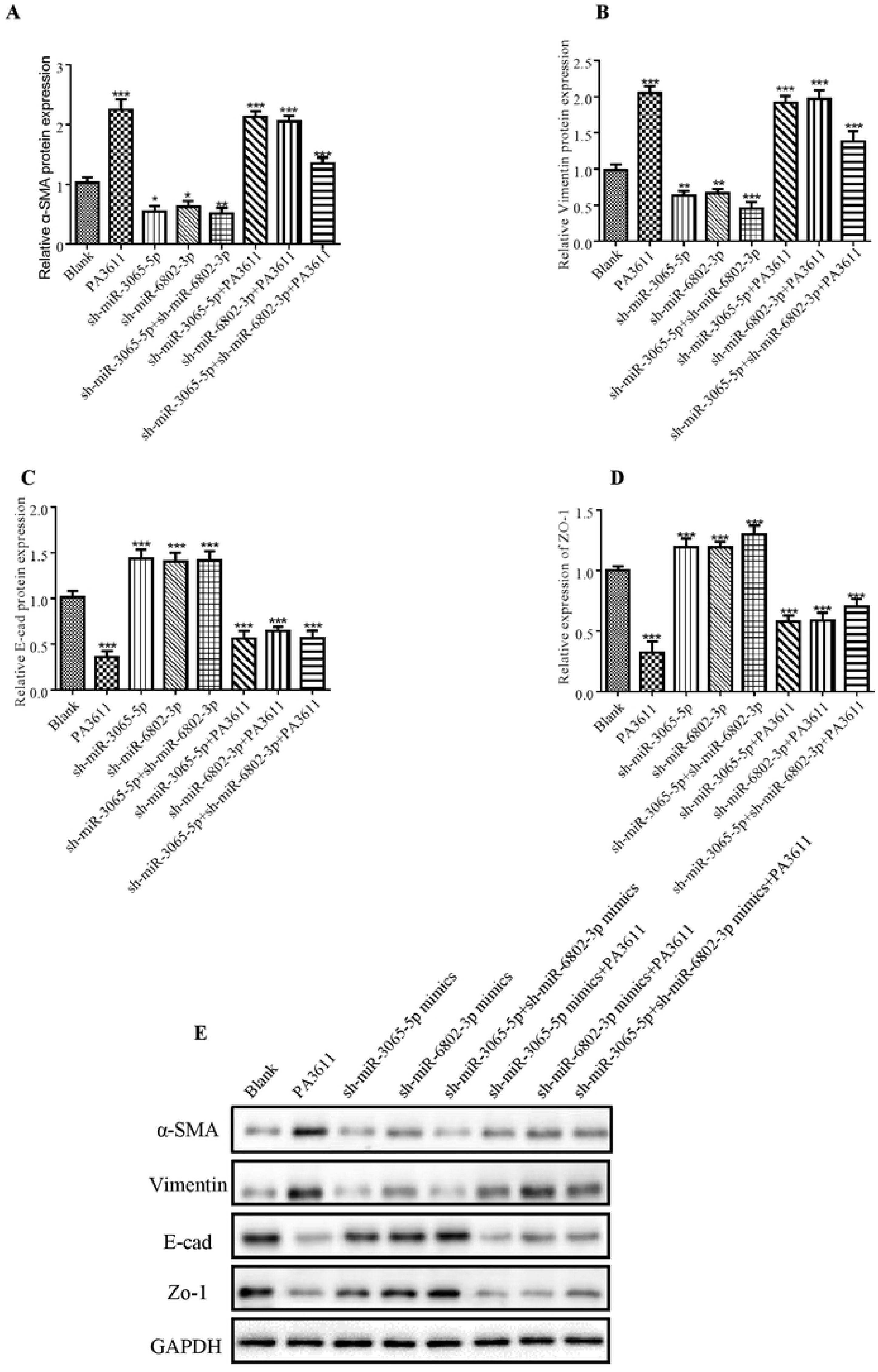
Effect of miR-3065-5p and miR-6802-3p knock down on the proteins expression of EMT relative markers. 16HBE cells were transfected with a negative control miRNA inhibitor (indicated with Blank), a miR-3065-5p inhibitor (indicated with sh-miR-3065-5p), a miR-6802-3p inhibitor (indicated with sh-miR-6802-3p), or both the miR-3065-5p and miR-6802-3p inhibitors (sh-miR-3065-5p+sh-miR-6802-3p). After 24h post transfection, cells were infected with PBS (negative control) or PA3611(30μg/ml) for another 24 hs, then cells were collected and detected for the proteins expression of EMT relative markers (Suppl Fig. 6A-E). The results were representative of three independent experiments. Data were presented as the means±SD. *, P<0.05, **, P<0.01, ***, P<0.001, vs. the control vector or the nonspecific siRNA interference group of the same treatment.

**Supplemental figure 7:**
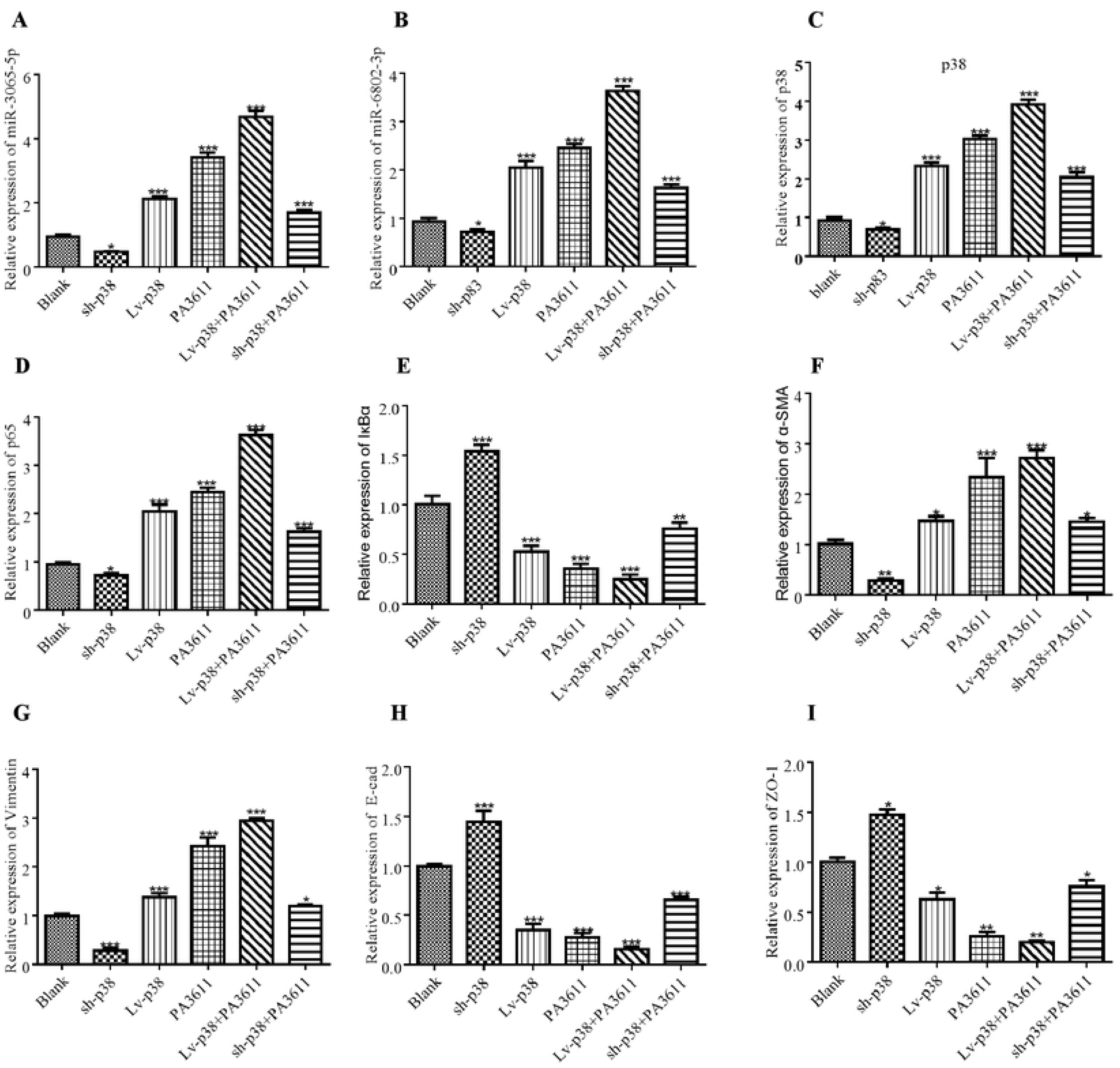
Effect of p38 intervention on the mRNAs expression of p38, IκBα, p65 and EMT relative markers. 16HBE cells were transfected with a control vector (indicated with Blank), a p38 overexpression vector (indicated with p38), or a specific siRNA directed against p38 (indicated with sh-p38). After 24h post transfection, cells were infected with PBS (negative control) or PA3611(30μg/ml) for another 24 hs, then cells were collected and detected for the mRNAs expression of p38, IκBα, p65 and EMT relative markers (Suppl Fig. 7A-I). The results were representative of three independent experiments. Data were presented as the means±SD. *, P<0.05, **, P<0.01, ***, P<0.001, vs. the control vector or the nonspecific siRNA interference group of the same treatment.

## Supporting information

**S1 Table.**
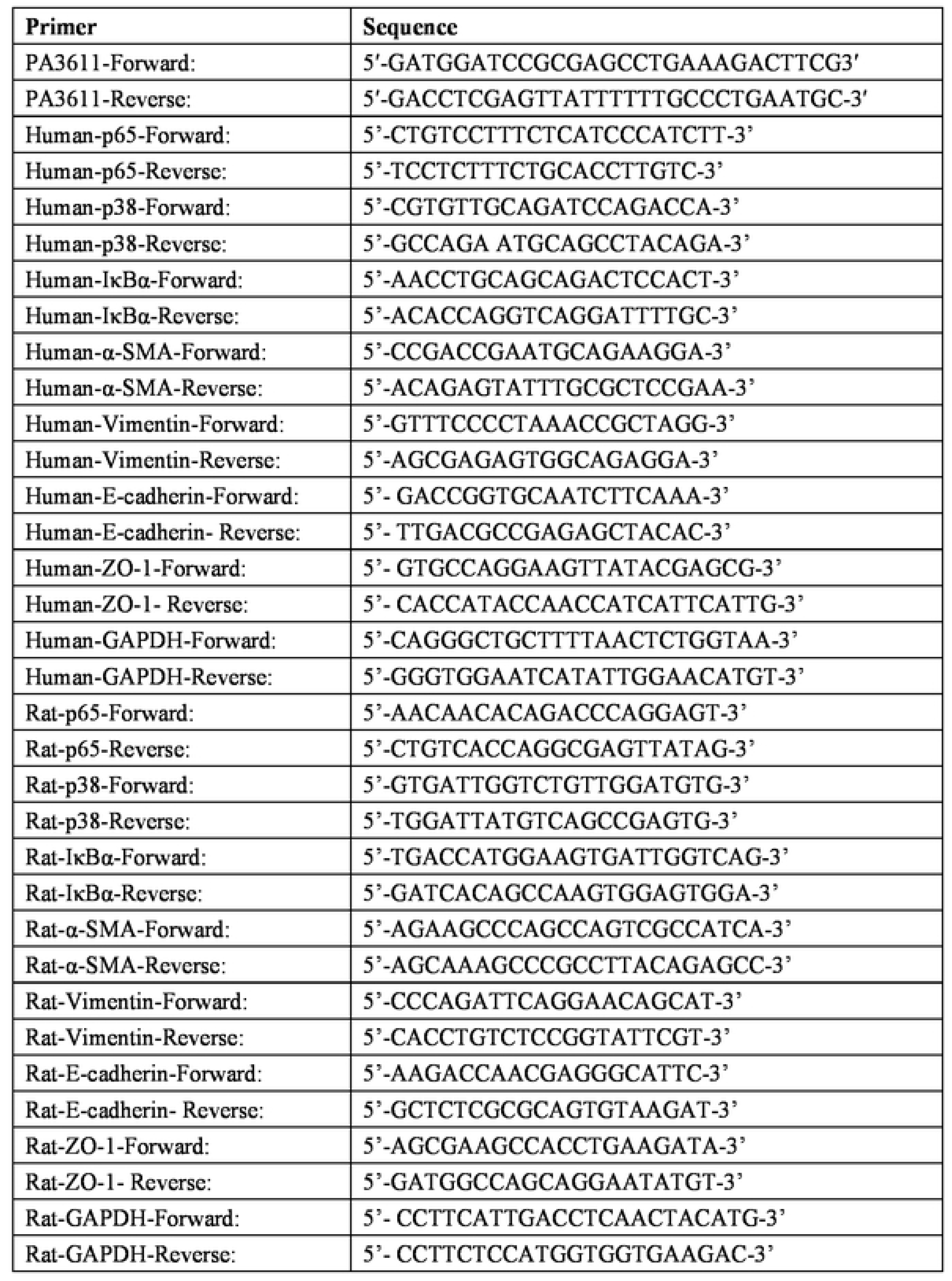
A list of primers used for PCR

**S2 Table.**
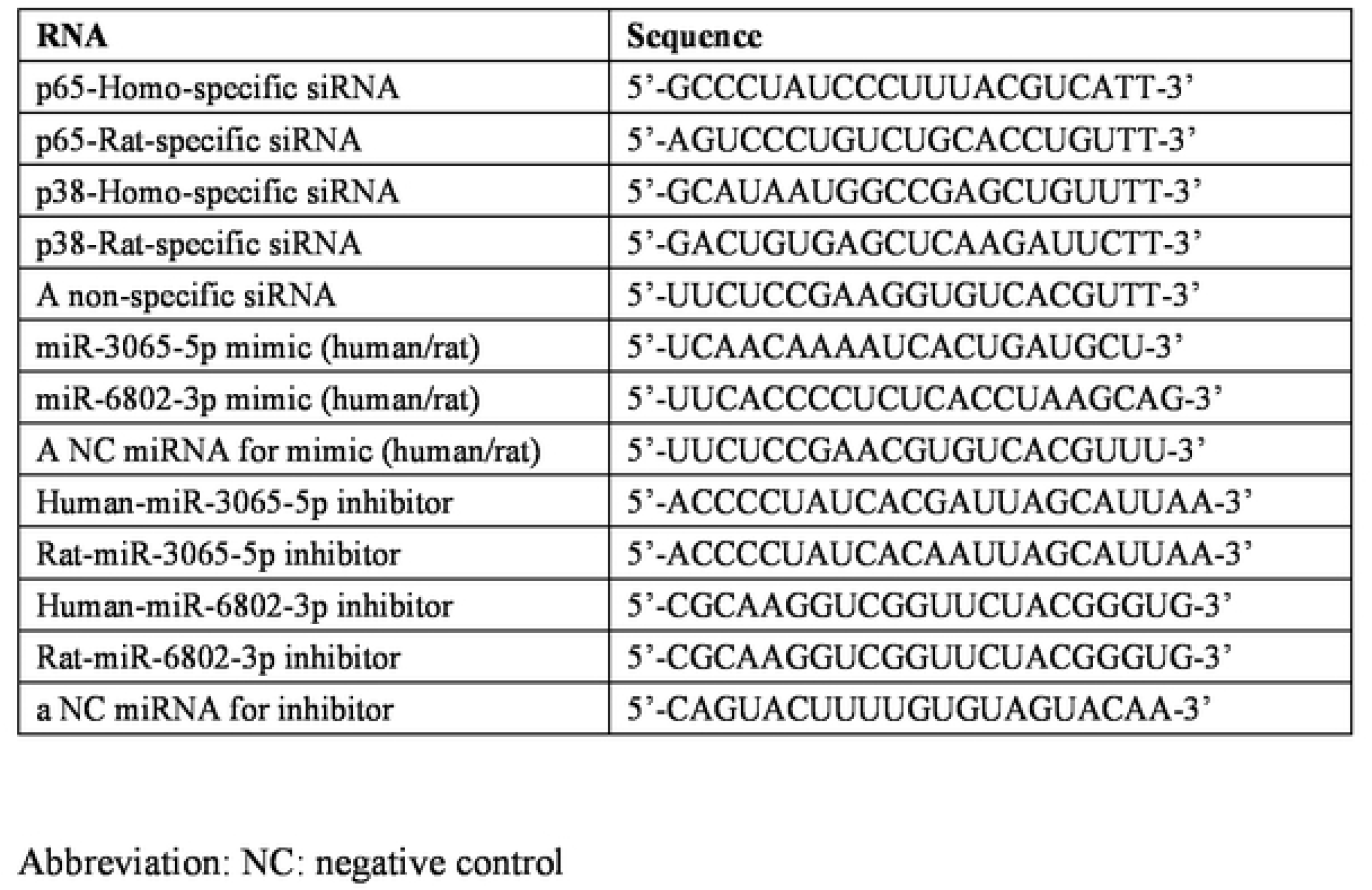
A list of RNA for transfection

